# Internal decay in living trees: a quantitative tomography framework and its application in a temperate forest

**DOI:** 10.64898/2026.06.10.730433

**Authors:** Grace Thompson, Maxwell P. Lutz, Taylor K. Lucey, Bethany Duncan, Masako Yang, Sam Jurado, Jaclyn Hatala Matthes, Robert E. Marra, Jonathan Gewirtzman

## Abstract

Internal decay in living trees is an important component of carbon and nutrient cycling as well as species and structural diversity maintenance in forest ecosystems. We used sonic and electrical resistance tomography to evaluate and compare the prevalence and severity of stem decay in 57 living trees among four common species (*Acer rubrum* L.*, Nyssa sylvatica* Marsh.*, Quercus rubra* L., and *Tsuga canadensis* (L.) Carrière)) with overlapping and non-overlapping distributions across wetland and upland habitat types at the Harvard Forest in Petersham, MA, USA. Independent of tree size, site identity best explained variation in the prevalence of decay across trees sampled, whereas species identity best explained the severity of decay. We categorized trees as having no decay, incipient decay, active decay, or cavities based on combined sonic and electrical resistance metrics, the latter generated by a custom image analysis application. About 31% of wetland trees exhibited incipient decay (compared to 11% in the upland), whereas about 32% of upland trees exhibited active decay (compared to 10% in the wetland). Our study highlights a new quantitative framework for decay categorization through normalized principal component analysis (PCA) and decay analysis software that complements dual tomographic methodology for future investigations of ecological drivers of decay presence and susceptibility.

## Introduction

Internal decay in living trees is a natural, fundamental process underlying forest community and ecosystem dynamics (Covey et al. 2012; Das et al. 2016; Oliva et al. 2020). Fungal pathogens and other microorganisms initiate decay by invading and decomposing either living or dead tissues, often through natural openings or injuries in tree shoot and root systems (Pearce 1996). Trees possess a repertoire of defense mechanisms that wall off, or compartmentalize, infections to prevent further internal proliferation (Shigo 1985). However, under favorable microsite conditions, wood decay fungi may break down those barriers, either directly or indirectly leading to host tree mortality by reducing resource acquisition, tree growth, and mechanical stability (Worrall et al. 2005; Oliva et al. 2012, 2014; González de Andrés et al. 2024). Indeed, trees with decay are more susceptible to snapping or uprooting due to the degradation of structural tissues (Larson and Franklin 2010). Resulting canopy gaps may release suppressed subcanopy tree species, simultaneously supporting biodiversity and structural complexity (Lewis and Lindgren 1999; Worrall et al. 2005; González de Andrés et al. 2024). Living trees with decay, standing dead trees, and downed logs provide habitat for decomposers and wildlife, as well as increase resource availability to the biological community (Franklin et al. 1987; Aitken and Martin 2007; Zuo et al. 2021). While internal decay is considered a background mortality agent and driver of deadwood accumulation (Franklin et al. 1987; Barrette et al. 2013; Wijas et al. 2024), its prevalence and extent remains understudied.

Decay occurrence and severity vary amongst tree species and individuals depending on a variety of factors specific to their wood properties, physiology, phenology, and ecology (Wagener and Davidson 1954; Morris et al. 2020; Kõrkjas et al. 2021). Some studies suggest that the likelihood of decay occurrence increases with tree age and diameter, as internal decay is a relatively slow process, and older trees tend to accumulate more wounds over their lifetimes (Frank et al. 2018; Kõrkjas et al. 2021; Gilbert et al. 2025). Additionally, tree species with slow life history strategies, characterized by denser wood, lower mortality rates, and better biochemical defenses, are considered less vulnerable to infection, decay, and mechanical instability compared to species with fast life history strategies (Chave et al. 2009). Ultimately, the ability of a tree to maintain a rate of increment growth that outpaces the spread of infection minimizes the likelihood that the decay column reaches the lateral vascular cambium (Shigo 1985; Morris et al. 2020).

Growth-defense tradeoffs may be further mediated by natural disturbances such as forest pests and pathogens that physically degrade wood and resources of tree hosts, potentially facilitating decay processes (Hicke et al. 2012; Dietze and Matthes 2014). Such natural disturbances are projected to increase across North America in response to more extreme climatic conditions (Hicke et al. 2012). Considering these observations, intra-and interspecific differences in tree longevity and decay resistance remain critical knowledge gaps (Di Filippo et al. 2015; Piovesan and Biondi 2021).

Existing estimates of decay prevalence and severity are limited geographically and methodologically (Johnstone et al. 2010; Frank et al. 2018), which has consequences not only for our understanding of forest ecological processes, but also of forest carbon dynamics. Live biomass accounts for a substantial proportion of carbon stocks in temperate forests (Pan et al. 2024) and overlooking internal decay may bias estimates of carbon storage and sequestration capacity (Marra et al. 2018). Many studies evaluating internal decay utilize tree felling or invasive survey techniques, such as resistance drilling and core sampling (Barrette et al. 2013; Heineman et al. 2015; Frank et al. 2018), while others are concentrated in forest management or urban settings (Ostrovský et al. 2017; Karlinasari et al. 2018; Brazee and Burcham 2023). Observations of wood decay fungal infections that lead to internal decay in living trees, characterized by discolored or decomposed wood, are often excluded from ground-based forest inventories (Zuleta et al. 2023). Further, external signs of decay, such as conks, tend to be inaccurate indicators of internal tree conditions (Marra et al. 2018; Gilbert et al. 2025). Therefore, gaps remain in the documentation of decay patterns in natural settings and across landscape gradients, as well as in understanding the consequences of these patterns for ecology and carbon.

To better understand the prevalence of internal decay and potential patterns within forests, more accessible, accurate, and less destructive methodologies must be validated and implemented (Soge et al. 2021). A growing body of evidence supports the use of a combination of minimally invasive tomographic methods to detect and characterize internal decay in standing, living trees (Brazee et al. 2011; Marra et al. 2018; Yue et al. 2019). For example, sonic tomography (SoT) and electrical resistance tomography (ERT) are designed to detect different wood physical properties, producing complementary images of tree cross-sections for more accurate assessments of the location, shape, extent, and stage of decay (Göcke et al. 2008). While tomographic methods are imperfect (Johnstone et al. 2010; Burcham et al. 2023), the repeatability and efficiency may provide essential supplementary information that strengthen projections of internal decay dynamics, from individual trees to forest communities.

We used SoT and ERT to document the prevalence and severity of internal decay within living trees at the Harvard Forest in Petersham, MA, USA. We created, validated, and implemented a custom image analysis application to calculate quantitative moisture-accumulation metrics from electrical resistance tomograms. Combining these results with sonic tomograms, we categorized the internal conditions of individual trees into four major categories: no decay, incipient decay, active decay, and active decay with a cavity. To evaluate the influence of environmental factors on decay patterns, we compared two species found in both wetland and upland habitats (red maple, *Acer rubrum* L. and eastern hemlock, *Tsuga canadensis* (L.) Carrière), as well as one upland (northern red oak, *Quercus rubra* L.) and one wetland specialist (blackgum, *Nyssa sylvatica* Marsh.). Our results provide a conceptual and quantitative framework for assessing decay trajectories in living trees, and provide insights into the prevalence and extent of decay in a natural forest setting.

## Methods

### Study site and species

This study took place during the summer of 2024 in the Prospect Hill Tract (585 ha) of Harvard Forest (42.5°N, 72.2°W), a temperate forest and long-term ecological research site in north-central Massachusetts, USA (Plotkin et al. 2015). Elevations range from 220 m to 410 m above sea level (Plotkin et al. 2015). Soils are characterized as acidic, stony, glacial tills, and are moderately to well drained in the upland regions and poorly drained in the wetlands (Jenkins et al. 2008). Between 1991 through 2024, the mean annual temperature was 8.29°C, and the mean annual precipitation was 1195 mm and spread relatively evenly throughout the year (Boose and VanScoy 2025). However, temperature has been increasing, and extreme precipitation events have become more frequent in the northeastern United States (Plotkin et al. 2015; Jurado and Matthes 2025).

Located within the transition hardwood region of New England, dominant tree species include *Q. rubra*, *A. rubrum*, *T. canadensis*, and eastern white pine (*Pinus strobus* L.) (Plotkin et al. 2015). In the Black Gum Swamp (BGS, about 12 ha), a forested peatland within Prospect Hill Tract, other dominant tree species include *N. sylvatica* and spruce species (*Picea* spp.) (Anderson et al. 2003; Jenkins et al. 2008). BGS is situated 365 m above sea level and composed of multiple basins that range from 2 m to 6.5 m in depth, with the deepest sedimentary layer dating over 14,000 years old (Anderson et al. 2003). Due to extensive clearing for agricultural purposes, followed by abandonment of these fields and establishment of commercial forestry practices into the twentieth century, most of these forests are even-age and second growth (Jenkins et al. 2008). Stands are primarily 75-120 years old, with some *T. canadensis* found to be greater than 200 years old and *N. sylvatica* greater than 300 years old (Plotkin et al. 2015).

*A. rubrum* and *Q. rubra* are both fast growing, early-to mid-successional species, the former being moderately shade tolerant and the latter being intermediately shade tolerant (Burns and Honkala 1990). As slow growing and highly shade tolerant species, *N. sylvatica* and *T. canadensis* may persist in understories for hundreds of years until periods of growth release and species turnover later in succession (Di Filippo et al. 2015; Thomas and Orwig 2025). *A. rubrum* thrives under a broad range of site characteristics, from swamps to dry uplands, due in part to their root system adaptability: seedlings respond to various environments by developing root system structures according to specific soil conditions, including moisture content fluctuations, and this functional trait is maintained through maturity (Burns and Honkala 1990). Likewise, local populations of *N. sylvatica* are tolerant to and successful under flooded conditions, with specialized physiological and morphological strategies associated with their root systems (Keeley 1979, 1980; Abrams 2007). On the other hand, *Q. rubra* prefer deep, well-drained soils characteristic of uplands (Burns and Honkala 1990); and *T. canadensis* prefer cool, moist soils, but are often avoidant of swampy or very dry sites (Thomas and Orwig 2025).

### Sampling design

The sampling design originally included 60 trees for SoT and ERT. Three tree species represented each study site in the upland (Environmental Measurement Station, EMS) and wetland (BGS), and 10 trees were selected per species. Present at both sites, *A. rubrum* and *T. canadensis* were considered the habitat generalists. *N. sylvatica* and *Q. rubra* were considered wetland and upland specialists, respectively. Individual trees were not selected based on signs of external decay. We used diameter at breast height (DBH, about 1.37 m from the ground) measurements from the Harvard Forest ForestGEO plot and selected trees that fell between the second and third quartiles of their respective species distribution (Fig. S1), disregarding any external decay indicators.

### Tomographic methodology

We used the PiCUS suite of tomographic equipment (Argus Electronic GmbH, Rostock, Germany), which included the PiCUS 3 Sonic Tomograph for SoT, the TreeTronic 3 Tomograph for ERT, and the Caliper 3 Geometry Measurement System. All tomographic measurements were conducted at DBH. For each tree, we placed a minimum of 8 sensors around the trunk circumference, following the manufacturer’s recommended sensor spacing based on trunk circumference. Sterilized steel nails were inserted through the bark to make solid contact with the wood, and sensor positions were recorded using the PiCUS Caliper 3, which determines cross-sectional geometry through triangulation measurements between sensor points.

#### Sonic tomography data acquisition and processing

SoT detects internal decay by measuring the velocity of acoustic waves traveling through wood (Brazee et al. 2011; Gilbert et al. 2016). Acoustic velocity is governed by the relationship between the modulus of elasticity (MOE) and density of wood; because decay reduces both MOE and density (depending on decay type), decayed and cavitated wood transmits sound more slowly than sound wood (Brazee et al. 2011; IML Electronic 2017). For SoT measurements, sensors were attached to the nails and connected to the PiCUS 3 Sonic Tomograph. Each measurement point was struck 3–5 times with the electronic hammer, and the system recorded the travel times of acoustic waves between all sensor pairs. Sound waves traveling through decayed regions or cavities take longer to traverse the cross-section than waves traveling through nondecayed wood; the PiCUS system compares measured travel times to expected travel times through solid wood within the same trunk rather than to an external reference, facilitating comparisons across species with different wood properties (Gilbert et al. 2016).

The PiCUS Q74 software uses a tomographic reconstruction algorithm to generate cross-sectional images displaying spatial variation in apparent sonic velocity. The resulting tomograms are displayed with a color scale where brown indicates high sonic velocity (sound wood), green indicates intermediate velocity (reduced density), and blue/magenta/white indicate low velocity (advanced decay or cavity). The software calculates the percentage of cross-sectional area in each velocity category based on the proportion of pixels in each color range. We exported the software-derived percent sound wood (brown) and percent damage (non-brown colors combined) for subsequent analysis. We did not perform independent image analysis of the sonic tomograms, instead using these software-derived percentages directly; for a detailed protocol and discussion of the considerations involved in image-based quantification of decay from sonic tomograms, we refer the reader to Gilbert et al. (2016). Previous validation studies have shown that, on average, SoT estimates of percent of decay are about 90% accurate when compared to visual estimates of felled stem disks, with discrepancies mostly consisting of underestimations by SoT (Gilbert and Smiley 2004; Johnstone et al. 2010).

#### Electrical resistance tomography data acquisition and processing

ERT provides complementary information about internal wood condition by measuring the spatial distribution of electrical resistance across the cross-section (Brazee et al. 2011; IML Electronic 2017). Electrical resistance in wood is influenced primarily by moisture content, ion concentration, and cell structure (IML Electronic 2017). Low resistance indicates elevated moisture content or ion concentration, conditions often associated with active fungal decay or bacterial wetwood, whereas high resistance indicates dry wood or cavities.

For ERT measurements, we removed the acoustic sensors and attached TreeTronic electrode clamps to the same nails used for SoT. The TreeTronic system sequentially injects current between electrode pairs and measures the resulting voltage distribution at all other electrodes. The spatial pattern of electrical resistance is reconstructed using an inverse algorithm and displayed as a two-dimensional tomogram with resistivity values in Ohm-m (IML Electronic 2017). In the standard color scheme, blue indicates low resistance (high moisture/conductivity), green and yellow indicate intermediate resistance, and red indicates high resistance (low moisture). Interpretation requires considering that tree species exhibit characteristic baseline resistance distributions; for example, temperate hardwoods that do not produce true heartwood (including *Acer* spp.) exhibit patterns with lower resistance in the sapwood and higher resistance in the heartwood (Brazee et al. 2011; IML Electronic 2017).

The PiCUS software represents resistivity on a color scale bounded by a user-defined minimum and maximum (default 30–1000 Ω·m). Because this range is set at the time of measurement and clips values beyond its bounds, it cannot be recovered afterward; appropriate bounds are species-and context-dependent and should be checked against the range of plausible values before scanning so that genuinely high or low resistivities are not saturated. For our temperate study species the default 30–1000 Ω·m range encompassed essentially all measured values, so clipping was negligible. We held the reconstruction options (smoothness and mesh fineness) at their software defaults and constant across all scans to ensure comparability (IML Electronic 2017).

Unlike the SoT software, the PiCUS software does not provide quantitative summary statistics for ERT tomograms, and prior studies have generally interpreted tomograms qualitatively. To extract numeric metrics from ERT images, we developed an open-source image analysis application in R (R Core Team 2025) using the Shiny framework (Chang et al. 2026) (Fig S2). The application extracts the colorbar from each exported tomogram image, constructs a log-scale calibration function mapping pixel colors to resistivity values based on user-specified calibration points, and delineates the cross-sectional boundary (either automatically by detecting the PiCUS measurement polygon, or manually) (Fig. 1). For each pixel within the boundary, the application assigns a calibrated resistivity value by matching its color to the nearest colorbar entry. Because the calibration is derived from each tomogram’s own colorbar, the application accepts images exported with the color scale normalized within each tomogram or mapped to the full fixed range; the two produce effectively identical reconstructed values, differing only negligibly through the software’s binning of colors.

**Figure 1.**
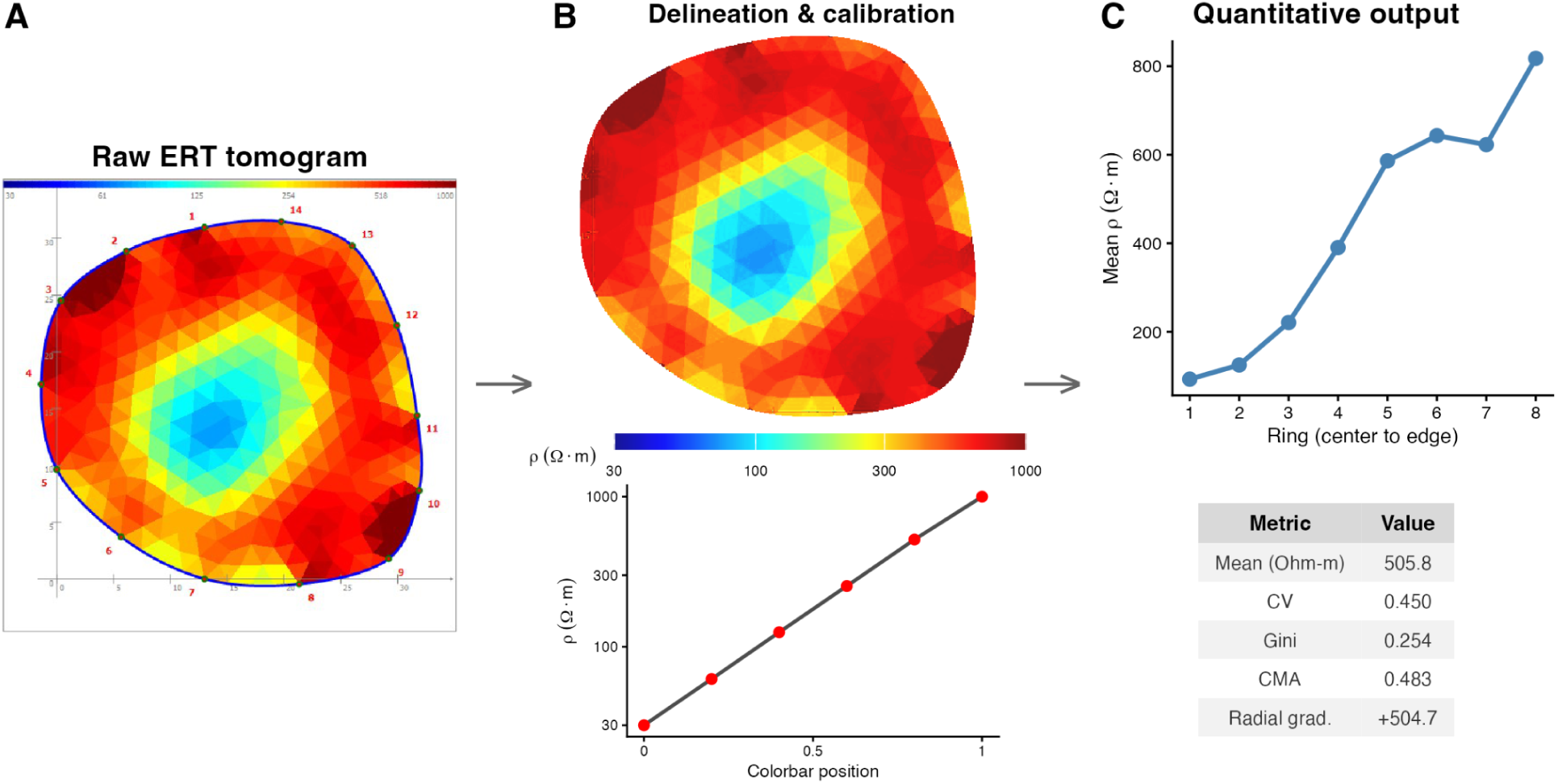
**Workflow for extracting quantitative resistivity metrics from PiCUS electrical resistance tomography (ERT) images**. (A) Raw ERT tomogram exported from PiCUS software showing the cross-sectional resistivity distribution of a *Q. rubra* individual with low to intermediate central resistance, or high central moisture accumulation; the colorbar (top) maps colors to resistivity values (30–1000 Ω·m) on a logarithmic scale. (B) Calibrated cross-section after automated boundary delineation and pixel-to-resistivity mapping, with the log-scale calibration function (lower panel; red points indicate user-specified calibration values). (C) Quantitative outputs: 8-ring radial profile of mean resistivity from the cross-section centroid to the edge (upper panel) and summary statistics including mean resistivity, coefficient of variation (CV), Gini coefficient, central moisture accumulation (CMA), and radial gradient (lower panel). The open-source analysis application is available at https://jgewirtzman-tree-tomography.share.connect.posit.cloud/ with source code at https://github.com/graceethompson/Tree-Tomography.

From the resulting resistivity map, the application computes the following metrics: mean and median resistivity (Ohm-m); standard deviation (SD); coefficient of variation (CV; SD/mean, a normalized measure of resistivity variability); Gini coefficient (a measure of inequality in the resistivity distribution, where 0 indicates uniform resistivity and higher values indicate concentration of extreme values); Shannon entropy (a measure of heterogeneity in the resistivity distribution); radial gradient (difference between mean resistivity in the outer third and inner third of the cross-section, where positive values indicate drier edges typical of healthy sapwood); and central moisture accumulation (CMA, the proportion of low-resistivity pixels (≤30th percentile) located in the inner third of the cross-section). CMA values near 0.33 indicate low-resistivity pixels are evenly distributed, while values substantially above 0.33 indicate anomalous moisture concentration in the heartwood. The application also generates 8-ring radial profiles of mean resistivity from the centroid to the edge.

#### Validation of ERT-moisture relationships

We selected 12 individuals of *T. canadensis* in the upland with available tree cores and three ERT images from below DBH, at DBH, and above DBH. These images were visually assessed to represent a potential gradient of decay. Tree cores were sampled using an increment borer at or slightly below DBH for the moisture content (%) of each tree core. Tree cores were weighed for their wet mass (g) and then kiln dried at 103 °C for 24 hours for their oven dry mass (g). Moisture content was estimated as the percentage of the mass of water in the wood (Glass and Zelinka 2021):

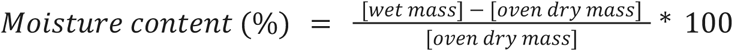

If the mass of water within a core was greater than the mass of the dry wood, then moisture content exceeded 100%.

### Decay classification system based on combined structural and moisture indicators

Unlike SoT, where sound wood produces a clear baseline of high and uniform velocity, the baseline resistivity distribution varies substantially among species, individuals, and seasons (Rust and Göcke 2008; IML Electronic 2017). To define the ERT axis, we standardized all eight ERT metrics from the application using species-specific z-scores and performed a principal component analysis (PCA) on the 57 main study trees. We used PC1 as a composite moisture anomaly index. Here we use “anomaly” in the sense of a signed deviation from a population central tendency, rather than to imply abnormality or pathology: because resistivity baselines vary by species, individual, and season, each metric was expressed as a departure from its species-specific mean, and PC1 is oriented so that higher values indicate lower resistivity (greater inferred moisture) and more heterogeneous moisture distributions than the species average. Trees were classified as having structural loss when SoT percent damage exceeded 1%, and as having anomalous moisture when their species-normalized PC1 exceeded the study-set mean.

We classified trees into four decay categories based on joint interpretation of SoT and ERT results, following criteria established by Brazee et al. (2011); Marra et al. (2018); and manufacturer guidelines (IML Electronic 2017) (Fig. 6): (A) No Decay—no structural loss and normal (high) resistivity patterns; (B) Incipient decay—no structural loss but anomalous resistivity (elevated moisture without measurable structural degradation); (C) Advanced (active) decay—structural loss with anomalous resistivity (moist, actively decaying wood); and (D) Cavity—structural loss with normal resistivity (hollow or desiccated decay column).

## Statistical analysis

One-way analysis of variance (ANOVA) was used to determine if the mean percent of damage detected by SoT was significantly different between sites. Two-way ANOVA was used to account for species-level differences to determine if species identity, site identity, or interactions between site and species had a significant effect on the percent of damage detected. If significance was detected, Tukey’s Honestly Significant Difference (HSD) post-hoc test was conducted for further pairwise comparisons.

Because the SoT data was zero-inflated (42 out of 57 trees (about 74%) exhibited no active decay or cavities), a hurdle modeling approach was implemented to model the occurrence, then severity of decay (Frank et al. 2018; Brazee and Burcham 2023). First, the probability of decay within each tree was modeled using binomial logistic regression, with decay presence (0/1) defined as the binary response variable. The explanatory variables considered were species (*A. rubrum*, *N. sylvatica*, *Q. rubra*, and *T. canadensis*) and site identity (BGS and EMS). A series of nested models, fit using the glm function in R, were built to test whether each explanatory variable significantly contributed to predicting the occurrence of decay, beyond what the null model or an antecedent explanatory variable could explain. Because not all species appeared in both sites, each explanatory variable was considered independent, or additive, with no interactions. Chi-square likelihood ratio tests were used to compare these models. To validate this binary approach, the fit of the full model with both variables was assessed using the simulateResiduals function in the DHARMa package in R (Hartig et al. 2024). Simulated residuals were uniformly distributed (KS test: p = 0.23), with neither over nor under dispersion (p = 0.816), and no outliers (p = 1).

Second, considering only the subset of trees with advanced decay (about 26% of trees), the severity of decay was modeled using beta regression, and incorporated the same explanatory variables from the first step. Because beta regression requires response values between 0 and 1, the raw percent damage values were converted to proportions prior to the analysis. Again, nested models were compared to test whether each explanatory variable significantly contributed to predicting the severity of decay within each tree. These models were fit with the betareg function (Zeileis et al. 2025). Goodness of fit was compared and assessed for each model using the Akaike information criterion (AIC). Model-predicted marginal probabilities and decay severity were estimated by averaging model predictions across covariate values. Bootstrap percentile confidence intervals (2,000 resamples) were computed by refitting each model to resampled datasets and re-averaging predictions.

## Results

The mean percent of damage detected by SoT across trees was 1.93% at the BGS site, which was significantly lower than the 7.07% detected at the EMS site (F(1, 55) = 5.143, *p* = 0.0273; Fig. 2). There also appeared to be less variability in the percent of damage among individuals at the BGS site compared to the EMS site (IQR equal to 0% versus 12%, respectively). However, further analysis indicated no significant species effect (F(3, 51) = 1.11, *p* = 0.354), site effect (F(1, 51) = 2.28, *p* = 0.137), or interaction effect of species and site (F(1, 51) = 0.16, *p* = 0.691) on the percent of damage. The complementary distribution of percent sound wood by site and species is shown in Fig. S3.

**Figure 2.**
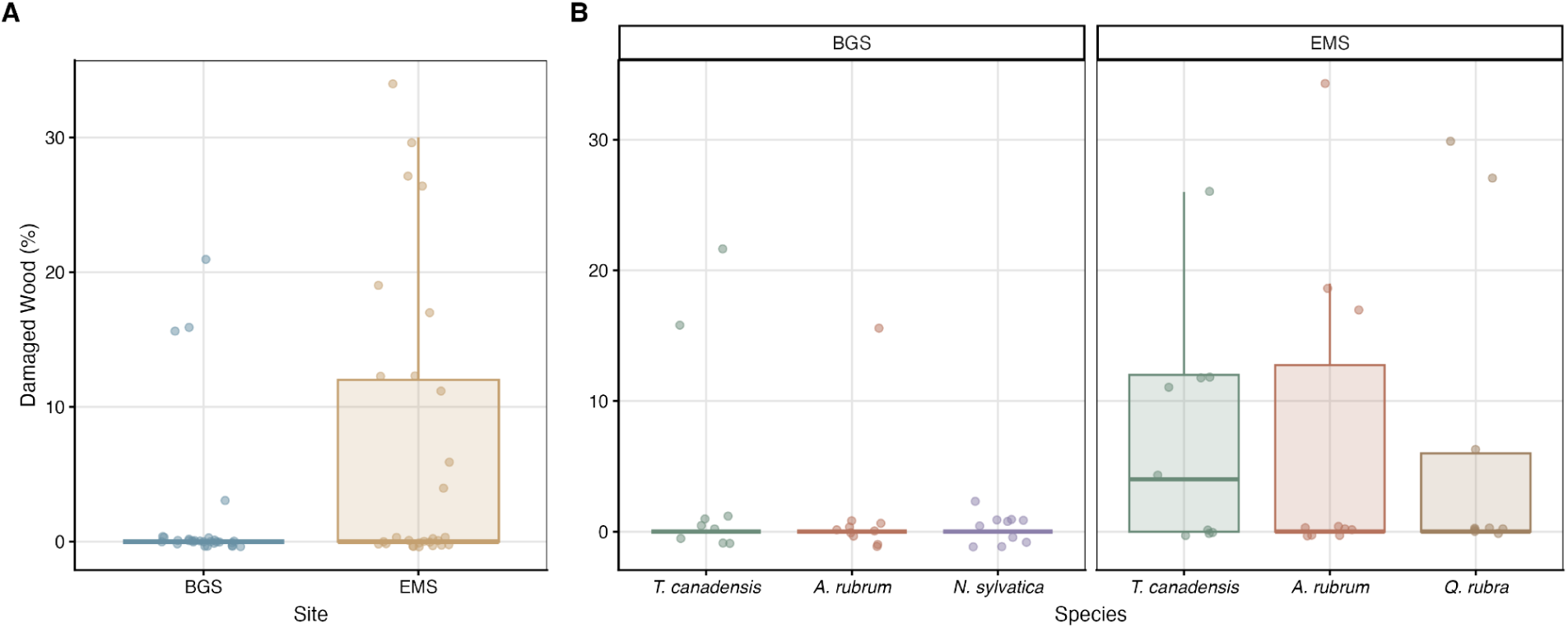
Boxplots of the percent of damaged wood for the BGS and EMS sites, for the tree species within each site. Jittered points represent raw data values for individual trees.

There was no significant site effect (F(1, 51) = 0.15, *p* = 0.704) or interaction effect of site and species (F(1, 51) = 0.04, *p* = 0.840) on mean electrical resistivity (Ω·m), but there was a significant species effect (F(3, 51) = 11.69, *p* < 0.001). Across sites, on average, *A. rubrum* (mean ± SE = 252 ± 26.2 Ω·m) had significantly lower resistivity than *T. canadensis* (mean ± SE = 381 ± 25.5 Ω·m, *p* = 0.0122745), *N. sylvatica* (mean ± SE = 447 ± 51.2 Ω·m, *p* < 0.001), and *Q. rubra* (mean ± SE = 519 ± 36.1 Ω·m, *p* < 0.001). *T. canadensis* also had significantly lower resistivity than *Q. rubra* (*p* = 0.0423252). Other species comparisons were not significantly different (*p* > 0.53). Resistivity heterogeneity, measured as the coefficient of variation (CV), also differed significantly among species (F(3, 52) = 4.57, *p* = 0.006) but not by site (F(1, 52) = 0.34, *p* = 0.561; Fig. 3C, D). *A. rubrum* exhibited the highest CV (mean ± SE = 0.655 ± 0.054), indicating the most heterogeneous resistivity distributions, and differed significantly from *T. canadensis* (0.459 ± 0.044, *p* = 0.024) and *Q. rubra* (0.397 ± 0.028, *p* = 0.015). Other species comparisons were not significant (*p* > 0.17). Median resistivity, SD, Gini coefficient, Shannon entropy, radial gradient, and CMA each showed significant species effects but no site effects (Figs. S4–S9).

**Figure 3.**
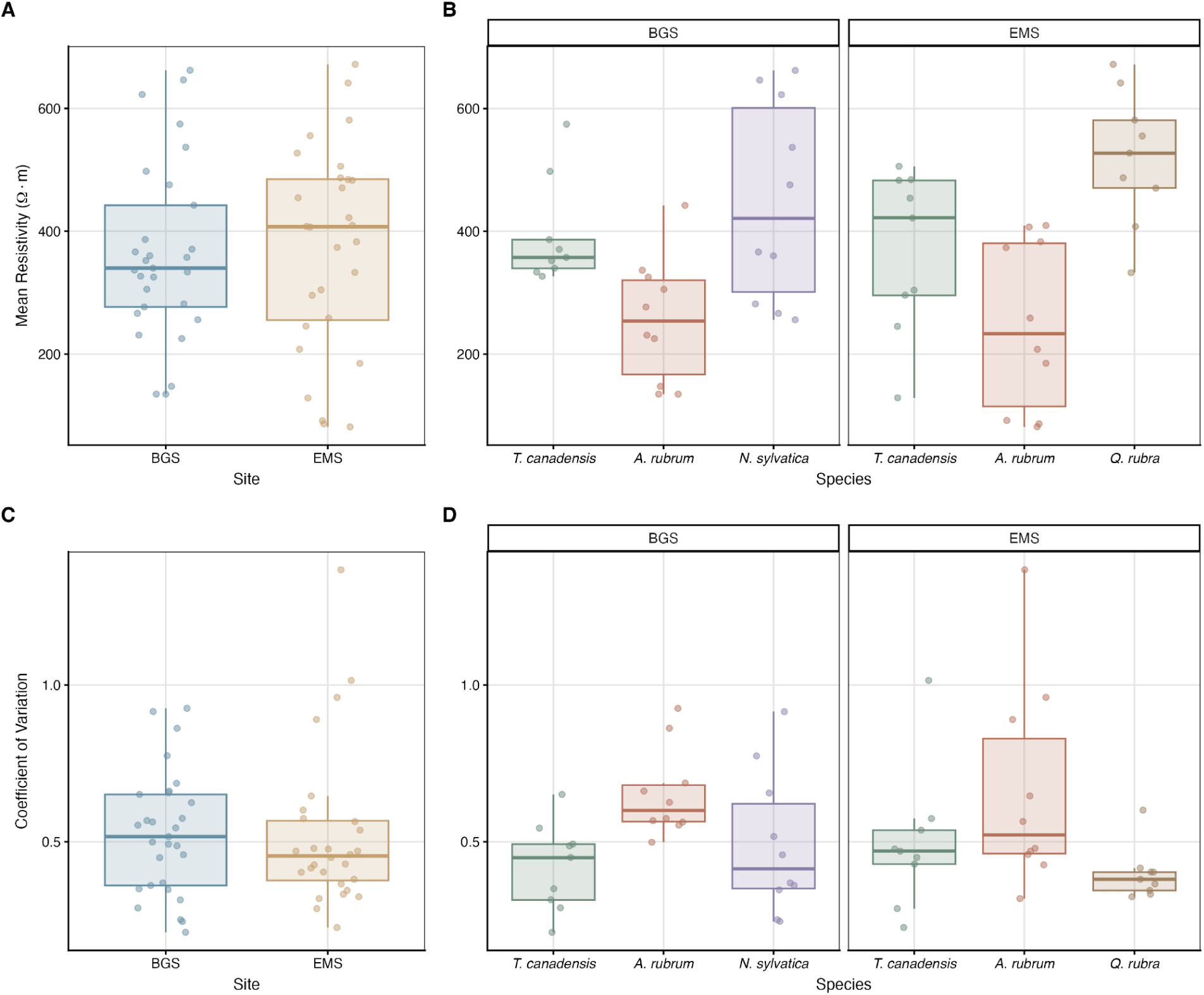
(A, B) Mean electrical resistivity (Ω·m) and (C, D) resistivity coefficient of variation for each species within the BGS and EMS sites. Jittered points represent raw data values computed by the open-source image analysis application for electrical resistance tomograms.

Site was the best predictor of decay prevalence (Fig. 4). The probability of decay presence differed between sites (χ^2^ = 4.91, df = 1, *p* = 0.02666), but not amongst species (χ^2^ = 3.67, df = 3, *p* = 0.29936). Adding species as a predictor did not improve model fit beyond site alone (*p* = 0.53082), but site did beyond species alone (*p* = 0.06332). Consistent with the full, additive binomial model, there were no significant species effect (*p* > 0.44), but there was a slightly positive site effect (β = 1.43 ± 0.81 SE, *z* = 1.77, *p* = 0.0772), suggesting a greater probability of decay at the EMS site compared to the BGS site.

**Figure 4.**
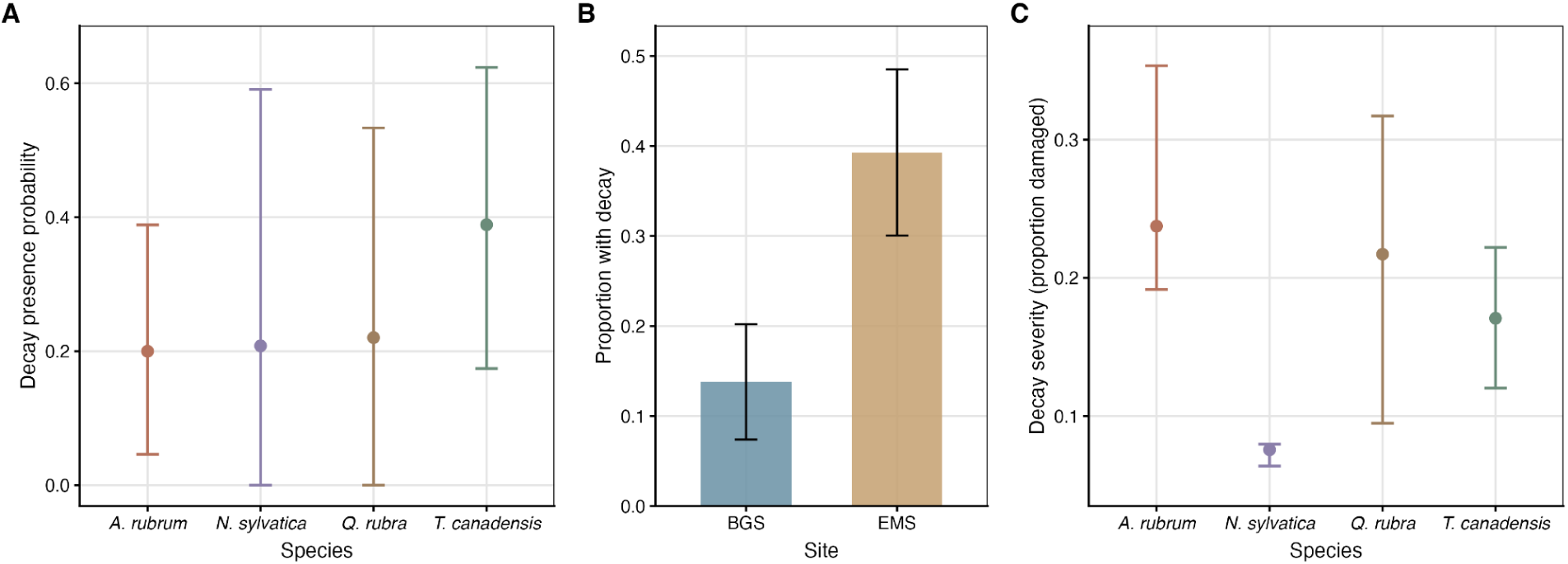
Hurdle model for decay prevalence and severity. (A) The probability of decay presence across species predicted by an additive binomial generalized linear model, with species and site as predictor variables. (B) Observed proportion of trees with decay at each site. Bars indicate the standard error. (C) Given that decay is present, decay severity across species predicted by a beta regression model, using only species as the predictor variable.

**Figure 5.**
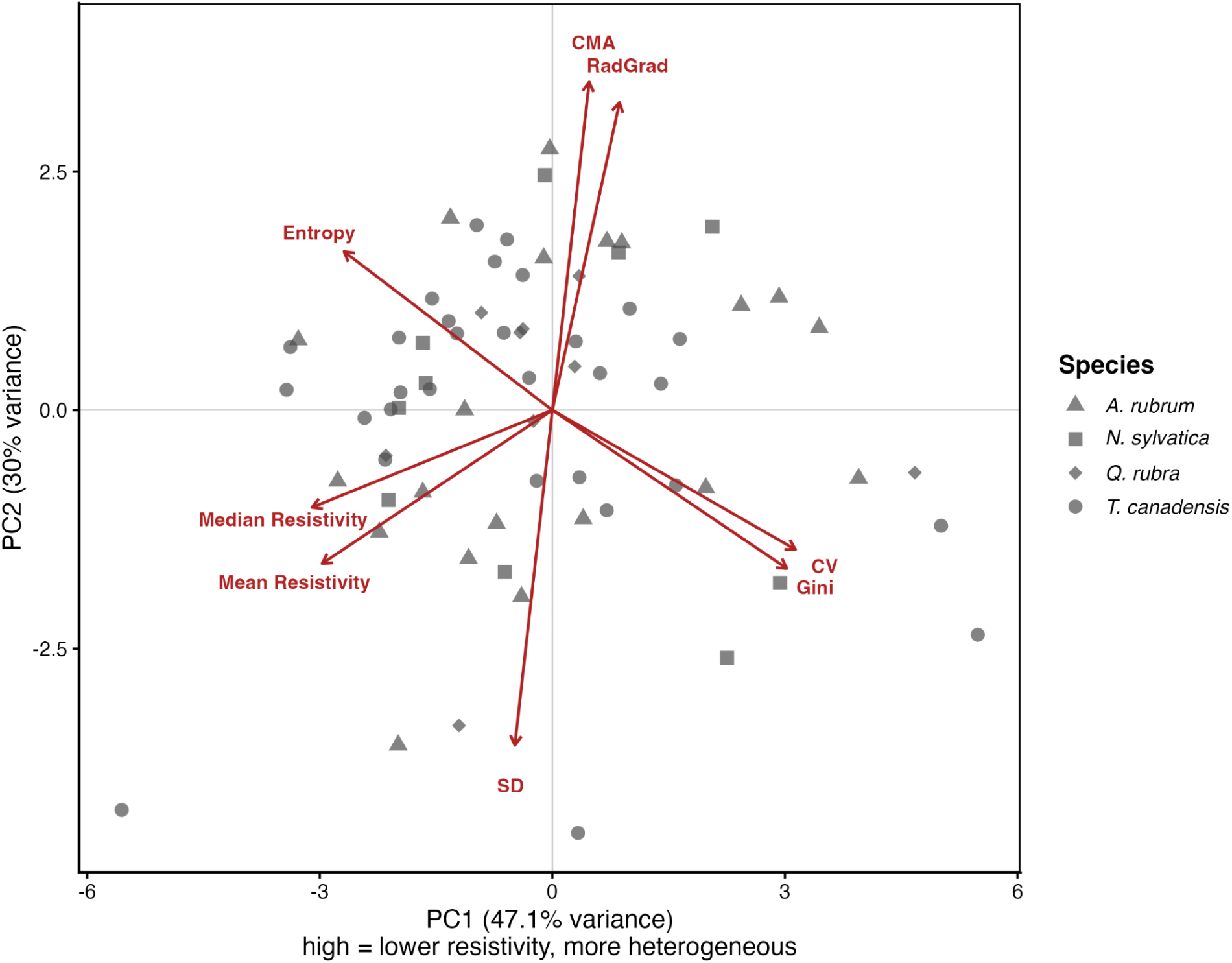
PCA biplot of eight species-normalized ERT metrics. PC1 (47.1% of variance) separates trees by metrics of central tendency (mean and median resistivity) and non-spatial variability (CV, Gini coefficient, entropy), where higher PC1 values indicate lower resistivity (i.e., wetter wood) with more heterogeneous moisture distributions. PC2 (30.0%) captures variation in spatially structured variability metrics (CMA, radial gradient). Note that resistivity is inversely related to moisture content, such that low mean/median resistivity corresponds to high wood moisture.

**Figure 6.**
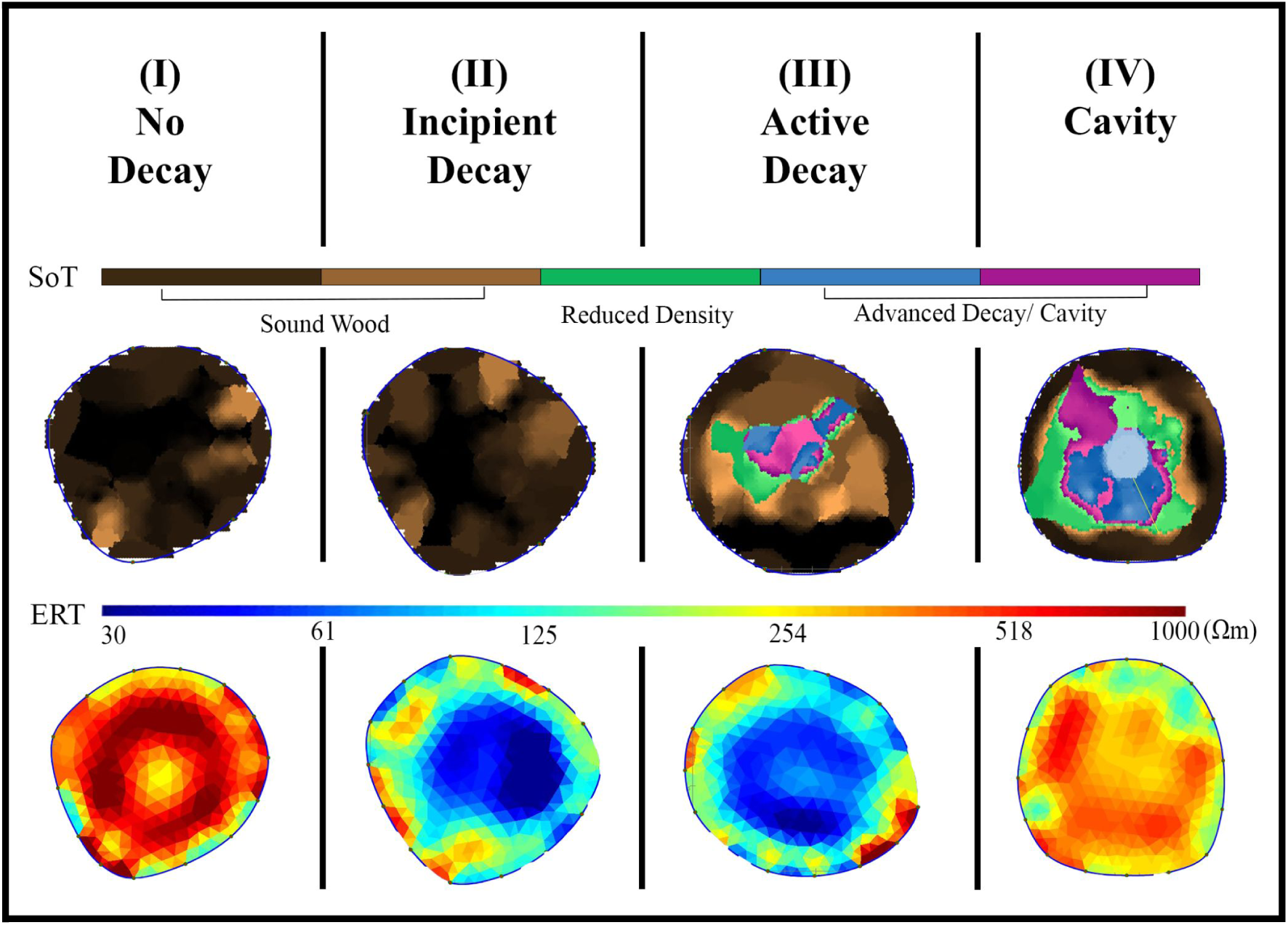
Representative sonic tomography (SoT) and electrical resistance tomography (ERT) cross-sections illustrating the four decay phases. Each column shows paired SoT (top) and ERT (bottom) tomograms from a single tree classified as (I) Sound, (II) Incipient Decay, (III) Active Decay, or (IV) Cavity. In SoT images, brown tones indicate sound wood and cooler tones (green, blue, violet) indicate reduced density or cavity formation. In ERT images, low resistivity (blue) corresponds to high moisture content and high resistivity (red/warm tones) corresponds to dry wood or cavity. Sound trees exhibit uniformly high density and high resistivity (low moisture). Incipient decay is characterized by intact structure (no SoT damage) with low resistivity, suggesting elevated internal moisture prior to detectable structural loss. Active decay shows both reduced density and low resistivity, consistent with concurrent structural degradation and elevated moisture. Cavity-stage trees display advanced structural loss with high resistivity in the decayed region, indicating desiccation of the remaining wood or air-filled voids. Resistivity color scale is logarithmic (30–1000 Ω·m).

Among trees with decay, species rather than site was the best predictor of decay severity (pseudo-R^2^ = 0.41). Including site as a predictor did not improve model fit (ΔAIC = 1.86). Compared to *N. sylvatica*, *A. rubrum* exhibited significantly greater decay severity (β = 1.70 ± 0.84 SE, *z* = 2.03, *p* = 0.042485), and *Q. rubra* slightly greater decay severity (β = 1.53 ± 0.85 SE, *z* = 1.79, *p* = 0.073896). *T. canadensis* decay severity was not significantly different compared to other species (*p* = 0.146).

We performed PCA on eight species-normalized ERT metrics to derive a composite moisture anomaly index. PC1 served as the ERT axis in our classification. PC1 explained 47.1% of total variance, with PC2 explaining an additional 30.0% (cumulative 77.1%). PC1 was driven by the contrast between distributional heterogeneity metrics (CV: +0.46; Gini: +0.45) and mean resistivity (−0.44), median resistivity (−0.46), and Shannon entropy (−0.40), such that higher PC1 values correspond to trees with lower overall resistivity and more heterogeneous moisture distributions. Spatial metrics (CMA: +0.07; radial gradient: +0.13) contributed modestly to PC1, loading primarily on PC2.

ERT PC1 correlated positively with gravimetric core moisture content across the 12 *T. canadensis* validation trees (moisture range: 67–142%; Pearson r = 0.68, 95% CI [0.18, 0.90], *p* = 0.015; Spearman ρ = 0.71, *p* = 0.009; R² = 0.46; Fig. S10). Mean resistivity alone correlated negatively with moisture (r = −0.72, *p* = 0.008), consistent with the known inverse relationship between resistivity and moisture content.

Of the 57 trees surveyed, 30 (53%) were classified as No Decay, 12 (21%) as in Incipient decay, 12 (21%) as in Active decay, and 3 (5%) as Cavity (Fig. 7). The majority of trees across all species were classified as Sound, ranging from 44% of *T. canadensis* to 60% of *N. sylvatica*. Incipient decay was most prevalent in *N. sylvatica* (30%) and *A. rubrum* (25%), suggesting elevated internal moisture without detectable structural degradation. Active decay was most common in *T. canadensis* (33%), the highest proportion among the four species surveyed. Cavity-stage trees were rare across all species (0–11%). Site-level differences were apparent: the upland EMS site had a higher proportion of trees with active decay (32%) compared to the wetland BGS site (10%), while BGS had a higher proportion of incipient decay (31% vs. 11% at EMS). The proportion of sound trees was similar between sites (55% at BGS, 50% at EMS).

**Figure 7.**
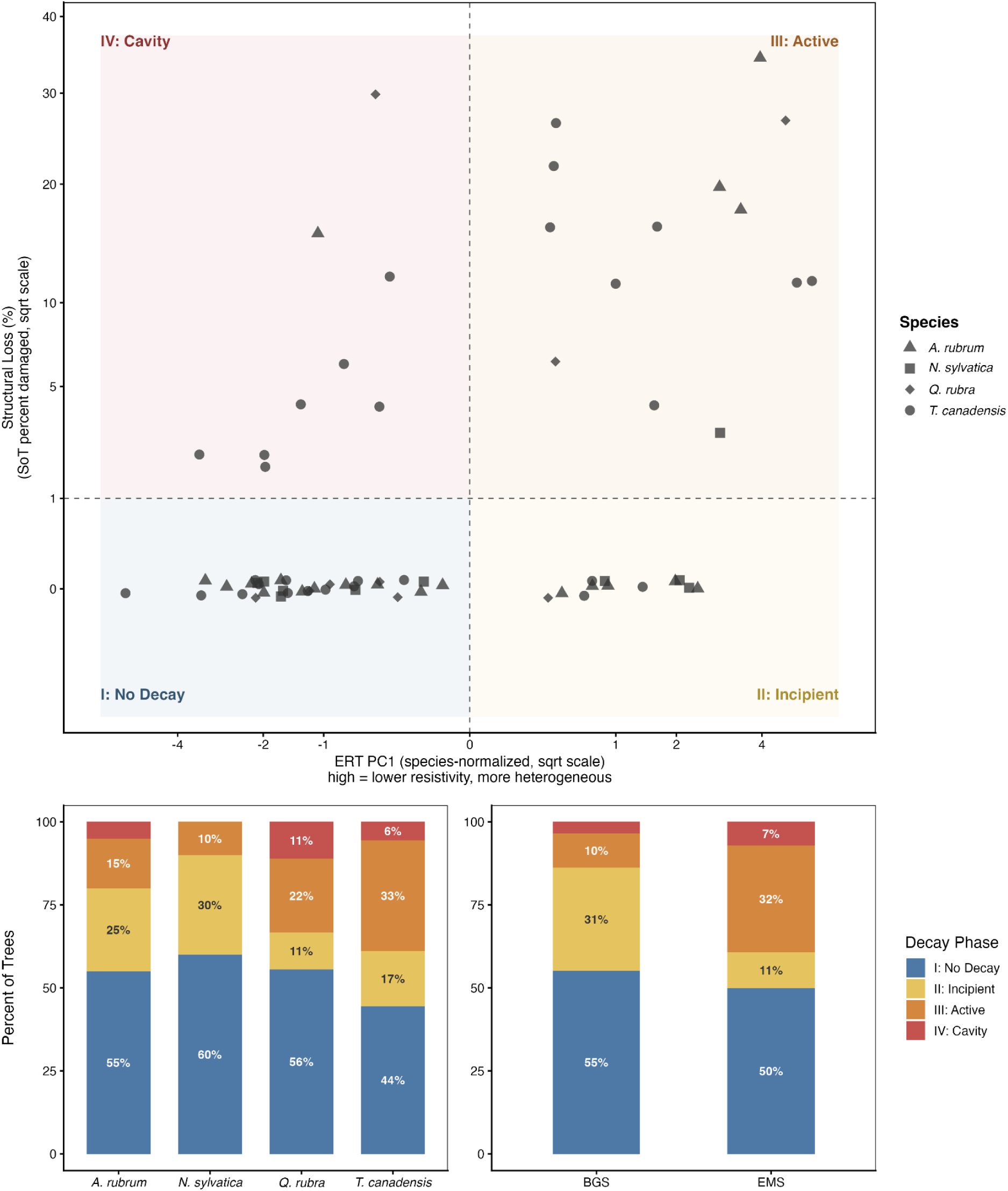
**Decay phase classification of 57 trees based on combined sonic tomography (SoT) and electrical resistance tomography (ERT) indicators**. Upper panel: two-dimensional phase diagram with SoT structural loss (percent damage, square-root scale) on the y-axis and species-normalized ERT PC1 (a composite moisture anomaly index; 47.1% of variance explained) on the x-axis. Dashed lines indicate classification thresholds (1% structural loss; study-set mean PC1). Trees are classified into four decay phases: I — No Decay (no structural loss, normal resistivity), II — Incipient decay (no structural loss, anomalous resistivity), III — Active decay (structural loss with anomalous resistivity), and IV — Cavity (structural loss with normal resistivity). Points are jittered slightly along the y-axis to reduce overplotting. Point shapes indicate species. Lower panels: relative frequency of decay phases by species (left) and site (right). BGS, Black Gum Swamp; EMS, Environmental Measurement Station.

## Discussion

We used SoT, ERT, and a novel image analysis application to both characterize and quantify internal decay within common tree species found in a wetland and upland region of a temperate forest. Our model predictions suggest that site conditions may be driving internal decay prevalence, while species-level traits may be driving severity. Trees in the upland forest had a greater proportion of internal decay compared to trees in the wetland. For trees with decay, both *A. rubrum* and *Q. rubra* exhibited greater decay severity compared to *N. sylvatica*. However, the hurdle model results were based on sonic tomograms alone, which detect structural loss but not early-stage moisture anomalies. In this way, any decay present was likely indicative of active decay or cavities. When combining data from the electrical resistance tomograms, we found more detailed differences in decay stages between sites and across species. For example, active decay was more prevalent amongst upland trees, whereas incipient decay was more prevalent amongst wetland trees. Documentation of similar patterns is limited along local soil moisture gradients in temperate forests, especially using minimally destructive tomographic methods (Frank et al. 2018; Marra et al. 2018; Aishan et al. 2024).

### Ecological drivers of decay prevalence and extent

Both within-and between-species variation in internal decay prevalence was best explained by site-level differences. While the probability of decay has been observed to increase with species’ flood tolerance and decrease with drought tolerance (Frank et al. 2018), (Hofmeyer et al. 2009) investigated the influence of soil drainage class on internal decay in three conifer species and, consistent with the overall trend we observed, found that decay incidence in northern white cedar (*Thuja occidentalis* L.) was greatest on well drained sites compared to poorly drained sites. Moisture availability in coarse woody debris is driven by soil moisture and wood characteristics, and tends to control the community composition and activity of decomposers, specifically enhancing decomposition by microorganisms, such as saprotrophic fungi (Chagnon et al. 2022; Wijas et al. 2024). Therefore, it would be reasonable to expect internal decay, especially advanced stages that parallel woody debris decomposition, to be most prevalent at our wetland site compared to our upland site. Yet excess moisture limits oxygen availability, which is an effective mechanism for living trees that prevents proliferation of wood decay fungi through functional sapwood, and underscores patterns of drier inner sapwood and heartwood layers being preferentially colonized (Boddy and Rayner 1983). Consistently, our ERT results showed that wetland trees had a higher proportion of incipient decay (31% at BGS vs. 11% at EMS), characterized by elevated internal moisture without structural loss, whereas upland trees more frequently exhibited active decay (32% at EMS vs. 10% at BGS). This pattern suggests that the anaerobic conditions of saturated sites may impede decay progression, maintaining elevated wood moisture without enabling the aerobic fungal activity required for structural degradation. In the better-drained upland soils, fungal colonization may proceed more readily from incipient moisture accumulation to active structural breakdown.

Notably, our sampling design controlled for the well-documented effect of tree size on decay probability by selecting individuals within the interquartile range of species-specific diameter distributions. The site and species effects we report are therefore assumed to be independent of diameter, the strongest known predictor of decay incidence given that decay occurrence tends to increase with age (Frank et al. 2018; Kõrkjas et al. 2021). Moreover, this constraint denotes that our results characterize decay patterns within the dominant canopy size class rather than across the full size spectrum, and future work spanning broader diameter ranges may reveal additional variation.

Natural and anthropogenic disturbances also influence internal decay prevalence (Shortle and Dudzik 2012). Windstorms, hydrological regime shifts, pest and pathogen infestations, and forest management represent the major disturbances observed at Harvard Forest (Plotkin et al. 2015; Jurado and Matthes 2025). Based on historic and contemporary records of these disturbances, BGS has remained largely unperturbed relative to upland regions in the Prospect Hill Tract (Jenkins et al. 2008). By way of windstorms and management practices, trees may acquire wounds, or wound other trees through branches breaking, tops snapping, partial uprooting, and tree felling, introducing multiple pathways for wood decay fungi to establish and having lasting effects on tree communities (Cline et al. 1991; Vasiliauskas 2001; Worrall et al. 2005). Large-scale rain events and drought, as well as pest and pathogen invasions, may simultaneously limit nutrient acquisition and tree growth, especially if trees have experienced some reduction in structural integrity (Dietze and Matthes 2014; Oliva et al. 2014; Vergara et al. 2026). Harsh soil conditions at BGS may limit the presence of pests and pathogens as well (Piovesan and Biondi 2021). These scenarios highlight the importance of complementing tomographic measurements with individual tree health and stand structure observations omitted from this study. Because BGS has experienced fewer wound-creating disturbance events than the upland (Jenkins et al. 2008), lower decay prevalence at BGS may partly reflect this differential disturbance legacy rather than, or in addition to, moisture-mediated suppression of fungal activity — a confounding result we cannot fully resolve with our design.

### Species-level variation in decay susceptibility and severity

While species identity was the strongest predictor of decay severity within trees, these findings were based solely on the proportion of decay detected by SoT, which does not accurately detect evidence of incipient decay (Brazee et al. 2011), such that the 26% of all trees sampled with decay detected were only either in advanced or cavity stages. Despite these limitations, interspecific differences suggest underlying variation in decay resistance among our focal species, aligning with expectations regarding taxonomic and functional group differences. Under the same edaphic condition, decay susceptibility is known to vary among hardwood and conifer species (Heineman et al. 2015; Piovesan and Biondi 2021).

*A. rubrum* has a poor ability to compartmentalize wounds and is considered highly susceptible to discoloration and decay, specifically by stem-rotting wood decay fungi (Burns and Honkala 1990). Additionally, as a short-lived and fast-growing subcanopy species, ecological factors such as competition and suppression may particularly reduce the overall vigor and development of *A. rubrum* individuals (Wagener and Davidson 1954). In contrast to *A. rubrum*, *Quercus* species form true heartwood with antimicrobial extractives that augment defenses against wood decay fungi (Pearce 1996). While decay resistance varies inconsistently across *Quercus* species, *Q. rubra* has been associated with a relatively high frequency of wood decay fungal pathogens and associated symptoms of decay (Brazee and Burcham 2023).

*N. sylvatica* and other tree species exhibit slower growth and increased investment in defenses in flooded and nutrient-poor soils, and these traits are related to decay resistance and longevity (Rodríguez-González et al. 2010; Di Filippo et al. 2015; Piovesan and Biondi 2021). Although *N. sylvatica* appeared to be the most resistant to decay, heart rot is characteristic of *N. sylvatica* and considered evident of this tradeoff between growth and defense (Abrams 2007; Piovesan and Biondi 2021).

Likewise, *T. canadensis* is considered a slow-growing species, resistant to decay (Thomas and Orwig 2025). However, hemlock woolly adelgid (*Adelges tsugae* Annand, HWA) depletes non-structural carbon stores that act as important energy sources for continued tree growth and survival (Thomas and Orwig 2025). Without adequate resources for defense, trees subjected to ongoing HWA infestation are likely more susceptible to the proliferation of decay fungi, which themselves deteriorate structural carbon (i.e., cellulose and lignin) (Shigo 1985). Notably, *T. canadensis* had the highest proportion of active decay (33%) among the four species despite its reputation for decay resistance, a pattern consistent with HWA-mediated weakening of defenses at this site. However, we did not assess HWA infestation status individually, so this link remains inferential.

### Internal moisture and the interpretation of electrical resistance tomography

ERT measures the spatial distribution of electrical resistivity across a stem cross-section, governed primarily by moisture content and ion concentration (Rust and Göcke 2008; IML Electronic 2017). Because actively decaying wood typically sustains elevated moisture as a consequence of fungal enzymatic activity and compromised cell wall integrity, ERT provides an indirect signal of biological processes that precede or accompany structural loss detectable by SoT (Brazee et al. 2011; Marra et al. 2018). However, this incipient fungal decay produces resistivity patterns essentially indistinguishable from bacterial wetwood (anaerobic, high-moisture heartwood) that does not itself degrade structural wood, an ambiguity inherent to the method (Sinclair and Lyon 2005; IML Electronic 2017; Marra et al. 2018). Wetwood may even suppress fungal colonization through oxygen exclusion (Worrall and Parmeter 1983), though this function can be lost if wounding aerates the colonized zone (Sinclair and Lyon 2005). Both *Tsuga* and *Acer* species, which lack strong decay-inhibitory heartwood extractives, are particularly prone to wetwood (Worrall and Parmeter 1983), making this ambiguity directly relevant to our sample. We follow (Marra et al. 2018) in designating Class B as “incipient decay or bacterial wetwood.”

Our normalized PCA approach is preferred to a single predefined metric because decay manifests differently across species: in some trees the signal is dominated by absolute moisture content; in others, by spatial heterogeneity. We validated ERT-derived metrics against wood moisture content from tree cores extracted from 12 *T. canadensis* individuals. ERT PC1 correlated positively with gravimetric core moisture content (r = 0.68, p = 0.015), confirming that the composite index tracks a real biological signal. Unlike a single core, the ability of PC1 to capture the spatial distribution and heterogeneity of moisture across the cross-section makes it sensitive to patterns of incipient decay that point sampling would miss. While limited in scope, this provides direct evidence that resistivity patterns correspond to measurable moisture differences, consistent with destructive validations by (Brazee et al. 2011) and (Marra et al. 2018). We acknowledge that ion concentration, temperature, and reaction wood can also influence resistivity (Rust and Göcke 2008), and that ERT captures a composite physicochemical signal rather than a direct measure of fungal activity. Its value lies in identifying anomalous moisture patterns that, combined with SoT, enable classification beyond what either method achieves alone (Deflorio et al. 2008; Brazee et al. 2011).

### Towards a quantitative framework for phased decay classification

The distinction between incipient and advanced decay is long established in forest pathology, describing a progression from early enzymatic degradation—where strength loss precedes weight loss—to structural breakdown and cavity formation (Hartig 1878; Cartwright and Findlay 1958; Boyce 1961; Wilcox 1978; Winandy and Morrell 1993). (Shigo and Sharon 1970) mapped this progression spatially as radial columns of discolored and decayed tissue emanating from wound sites, providing an anatomical basis for the radial moisture and density gradients captured by tomography. The CODIT model (Shigo 1985) describes host defense barriers rather than decay progression per se, and is complementary to the staged framework we employ. The four-category classification—No Decay, Incipient decay, Active decay, and Cavity—was formalized by Marra et al. (2018; see Table 1) as a 2×2 matrix crossing structural integrity (SoT) with moisture status (ERT), validated against 39 destructively sampled hardwood trees. The combined approach was motivated by the recognition that SoT alone cannot reliably detect incipient decay when structural integrity is preserved (Deflorio et al. 2008), and was developed collaboratively between researchers and instrument developers (Rust and Göcke 2008; Brazee et al. 2011; IML Electronic 2017).

Our study extends this framework from categorical to quantitative. We developed a processing pipeline that extracts continuous moisture metrics from ERT images, replacing subjective visual assessment with reproducible descriptors. We used PCA to derive a data-driven moisture anomaly threshold rather than relying on expert judgment. And by applying the framework across species and sites, we demonstrate that the four categories are ecologically interpretable, corresponding to different combinations of site conditions and species traits.

We interpret the four categories as diagnostic states, not as a validated temporal trajectory. Although the ordering “no decay → incipient → active → cavity” is conceptually consistent with progressive degradation (Boyce 1961; Schwarze et al. 2000), our cross-sectional design cannot confirm sequential transitions. Alternative pathways are plausible: incipient conditions may persist where oxygen limitation suppresses fungal progression (Rayner and Boddy 1988); trees may transition directly from sound to active decay; and cavities may represent stabilized endpoints. For example, tree species with relatively high longevity, such as *N. sylvatica*, often exhibit and maintain hollow stems for years (Di Filippo et al. 2015; Piovesan and Biondi 2021). Our classification also depends on operationally motivated thresholds—1% SoT damage and the study-set mean of PC1—and alternative thresholds could shift assignments near category boundaries.

Despite these caveats, the framework provides a structured, reproducible means of characterizing internal condition that goes beyond binary detection (Zuleta et al. 2023), capturing variation within the live biomass pool currently invisible to forest carbon accounting (Marra et al. 2018; Pan et al. 2024). While the 2×2 matrix cannot resolve incipient fungal colonization from bacterial wetwood, it nonetheless differentiates sound wood, early microbial colonization, active structural loss, and cavity formation—stages that carry distinct implications for tree stability and carbon storage. Nearly half (47%) of surveyed trees exhibited some form of internal anomalies detectable by tomography, suggesting that conventional allometric equations treating standing live biomass as intact may meaningfully overestimate carbon stocks at the stand level (Marra et al. 2018; Zuleta et al. 2023). Integrating tomographic assessments into inventory protocols represents a tractable path toward correcting this bias: time series measurements, where each of the decay stages are tracked over time to determine the probability that one stage leads to the next, may facilitate more accurate estimations of temporal variations in stored carbon.

## Conclusions

We documented and evaluated the four major stages of internal decay within *A. rubrum*, *N. sylvatica*, *Q. rubra*, and *T. canadensis*. Our study is one of the first to have explored internal decay patterns among these dominant tree species at Harvard Forest using dual tomographic methods alone. Moreover, we developed a novel application tool to analyze and supplement electrical resistance tomograms with quantitative summary statistics, reducing potential biases associated with visual resistivity assessments. Internal decay varied intraspecifically and interspecifically, and between wetland and upland environments. Specifically, we found that internal decay prevalence and severity were best explained by site and species identity, respectively. These results underline the importance of assessing ecological factors and functional traits that may influence decay susceptibility and resistance mechanisms within forest stands and different tree species. For more comprehensive evaluations of internal decay patterns within communities and their associations with ecosystem processes, such as carbon cycling and ecological succession, subsequent studies should maximize their tomographic observations, incorporate a broader range of tree diameters, and detail visible signs of injury and decay. Without destructive sampling, our study demonstrated the advantageous, accurate characterization and quantification of internal decay using SoT in tandem with ERT to support future, broader scale analyses.

## Acknowledgements

We thank Audrey Barker-Plotkin, David Orwig, and the Harvard Forest Woods Crew for field and logistical support, and the staff of the Harvard Forest Summer Research Program in Ecology. We thank Nicholas Brazee for guidance on tomographic methods. We thank Nicholas Ward for testing the ERT image-analysis application and for helpful discussion. This work was supported by the Harvard Forest Summer Research Program in Ecology (National Science Foundation REU grants DBI-1950364 and DBI-2348924). J.G. was supported by a National Science Foundation Graduate Research Fellowship (DGE-2139841), a Harvard Forest LTER Graduate Student Research Award, the Kohlberg-Donohoe Research Fellowship, and the Yale Institute for Biospheric Studies. Additional support was provided by the Department of Energy Environmental System Science grant DE-SC0024092 to J.H.M. The Harvard Forest Long-Term Ecological Research program (National Science Foundation grants DEB-8811764, DEB-9411975, DEB-0080592, DEB-0620443, DEB-1237491, DEB-1832210) provided site access and infrastructure.

## Supplemental Information

**Figure S1.**
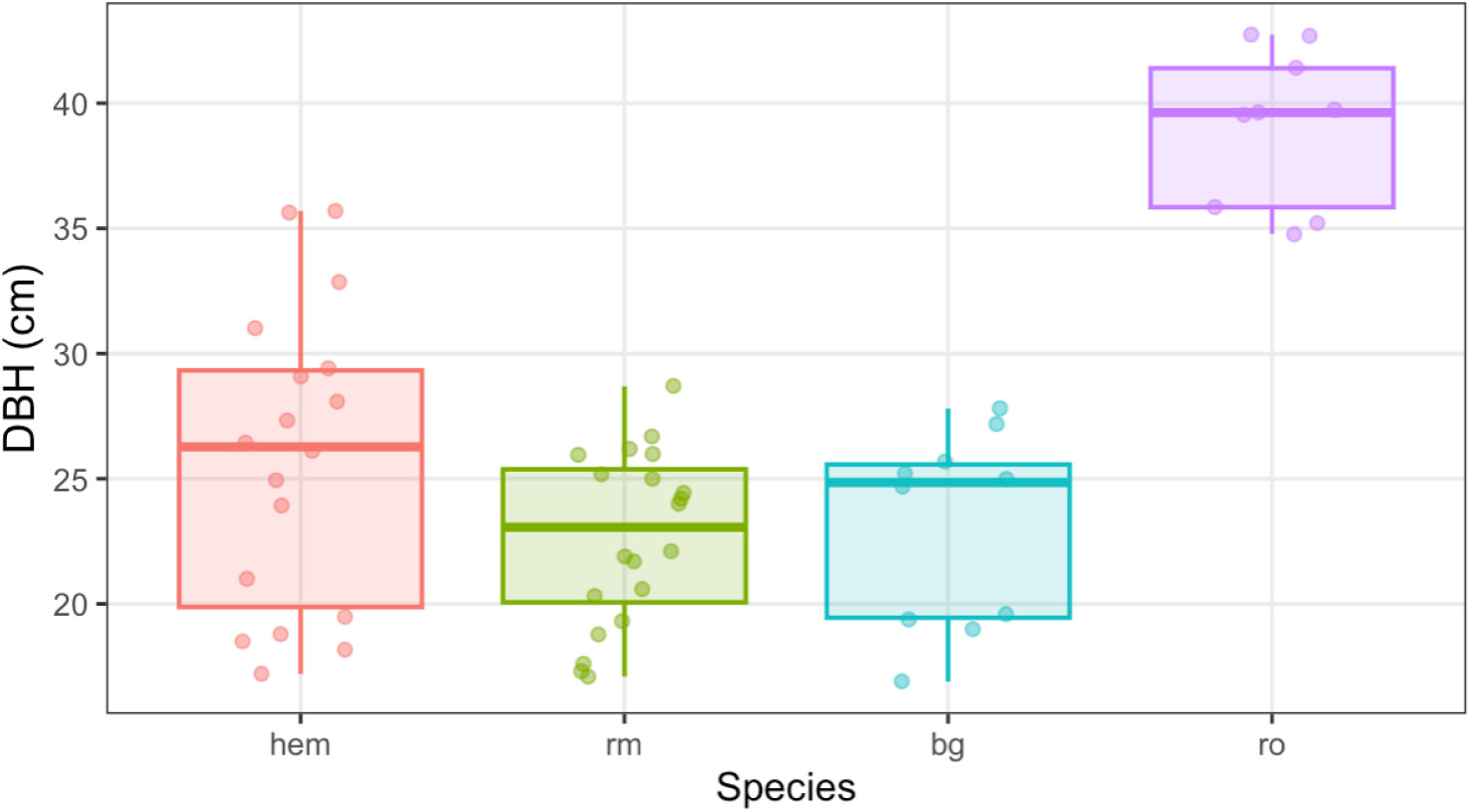
Diameter at breast height (DBH, cm) boxplots for each species. Jittered points represent raw data values for individual trees. Species differed significantly in mean DBH (one-way ANOVA: F(3, 53) = 31.05, p < 0.001). Q. rubra had the largest mean DBH (mean ± SE = 39.1 ± 1.03 cm), which was significantly greater than T. canadensis (25.8 ± 1.41 cm), A. rubrum (22.7 ± 0.78 cm), and N. sylvatica (23.0 ± 1.24 cm; all p < 0.05 by Tukey HSD). Other pairwise comparisons were not significant. Trees were selected within the second and third quartiles of each species-specific DBH distribution, so these values reflect the dominant canopy size class within each species rather than the full population range.

**Figure S2.**
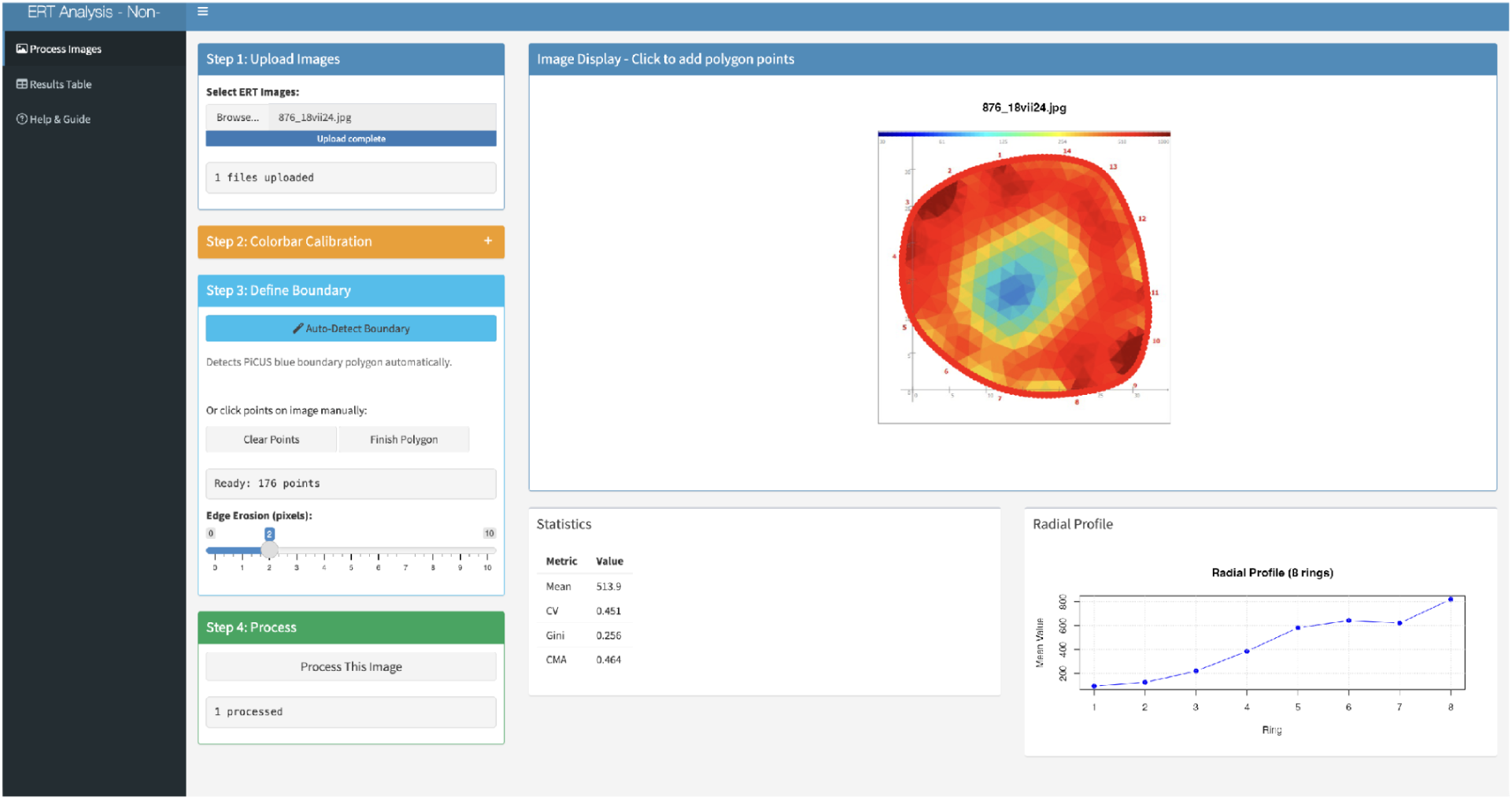
Screenshot of the open-source ERT image analysis application (R Shiny). The application extracts the colorbar from each exported PiCUS tomogram, constructs a log-scale pixel-to-resistivity calibration, delineates the cross-sectional boundary, and computes summary metrics. The application is available at https://jgewirtzman-tree-tomography.share.connect.posit.cloud/ with source code at https://github.com/graceethompson/Tree-Tomography.

**Figure S3.**
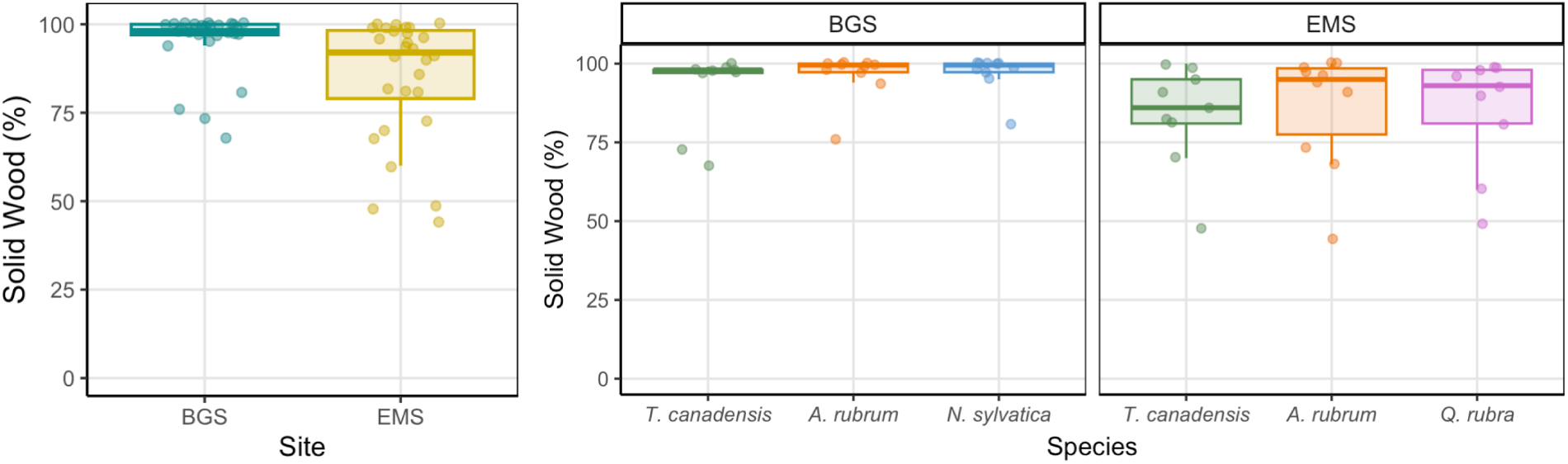
**Percent of cross-sectional area classified as sound wood by sonic tomography (SoT)**, shown as boxplots for each site and each species within sites. Jittered points represent raw data values for individual trees. Percent sound wood was significantly greater at BGS than EMS (Wilcoxon rank-sum test: W = 599, p = 0.002; two-way ANOVA site effect: F(1, 51) = 4.25, p = 0.044), reflecting the lower prevalence of structural decay at the wetland site. There was no significant effect of species (F(3, 51) = 1.42, p = 0.248) or species-by-site interaction (F(1, 51) = 0.04, p = 0.848).

**Figure S4.**
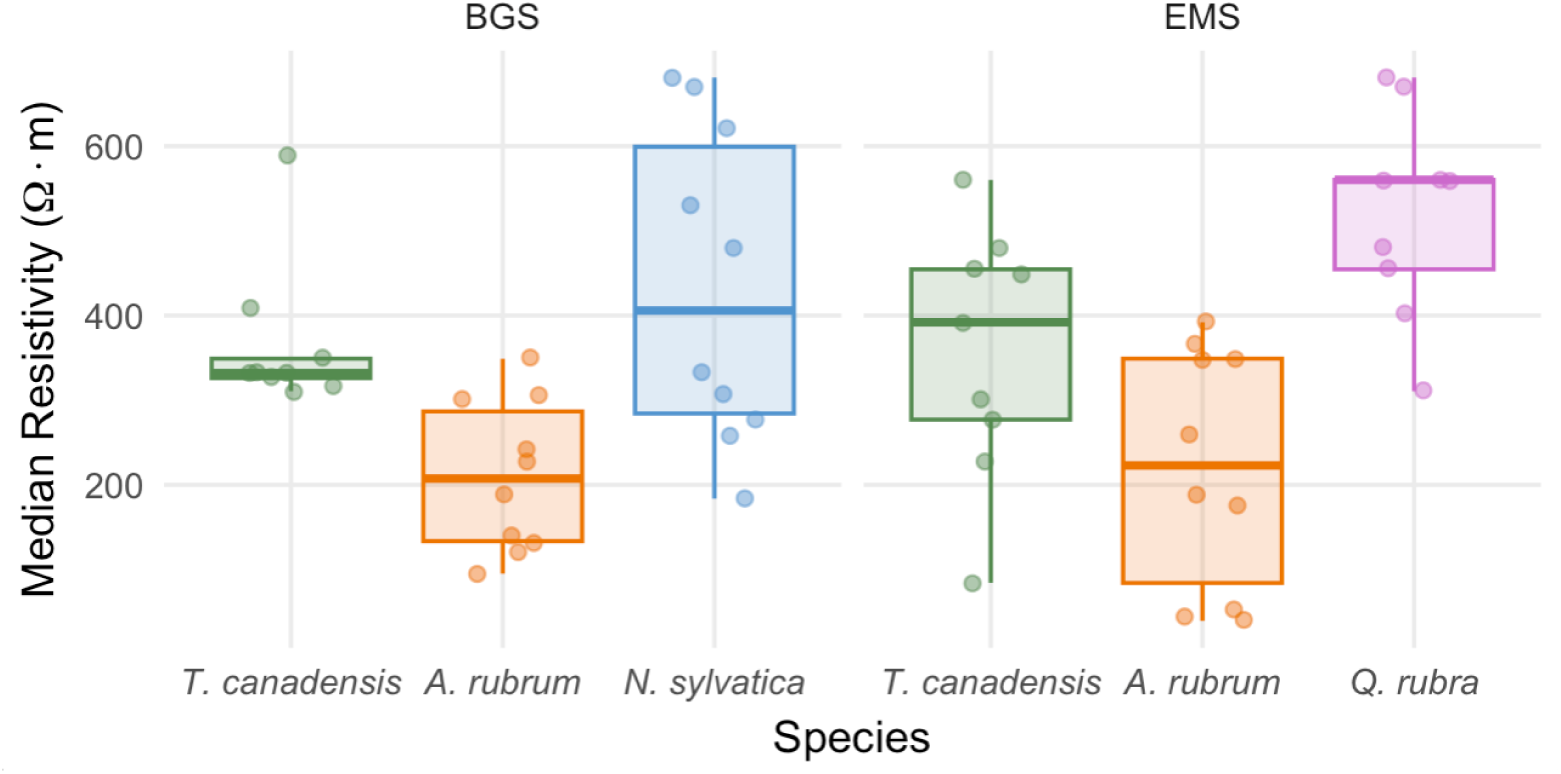
Median electrical resistivity (Ω·m) boxplots for each species within the BGS and EMS sites. Jittered points represent raw data values computed by the open-source image analysis application. Higher values indicate drier internal wood conditions. There was no significant site effect (F(1, 51) = 0.003, p = 0.958) or species-by-site interaction (F(1, 51) = 0.05, p = 0.824), but there was a significant species effect (F(3, 51) = 12.87, p < 0.001). Averaged across sites, median resistivity was significantly lower in A. rubrum (mean ± SE = 216 ± 25.8 Ω·m) than T. canadensis (362 ± 27.9 Ω·m, p = 0.008), N. sylvatica (434 ± 58.5 Ω·m, p < 0.001), and Q. rubra (520 ± 40.3 Ω·m, p < 0.001). Q. rubra also had significantly greater median resistivity than T. canadensis (p = 0.029). All other pairwise comparisons were not significant (p > 0.50).

**Figure S5.**
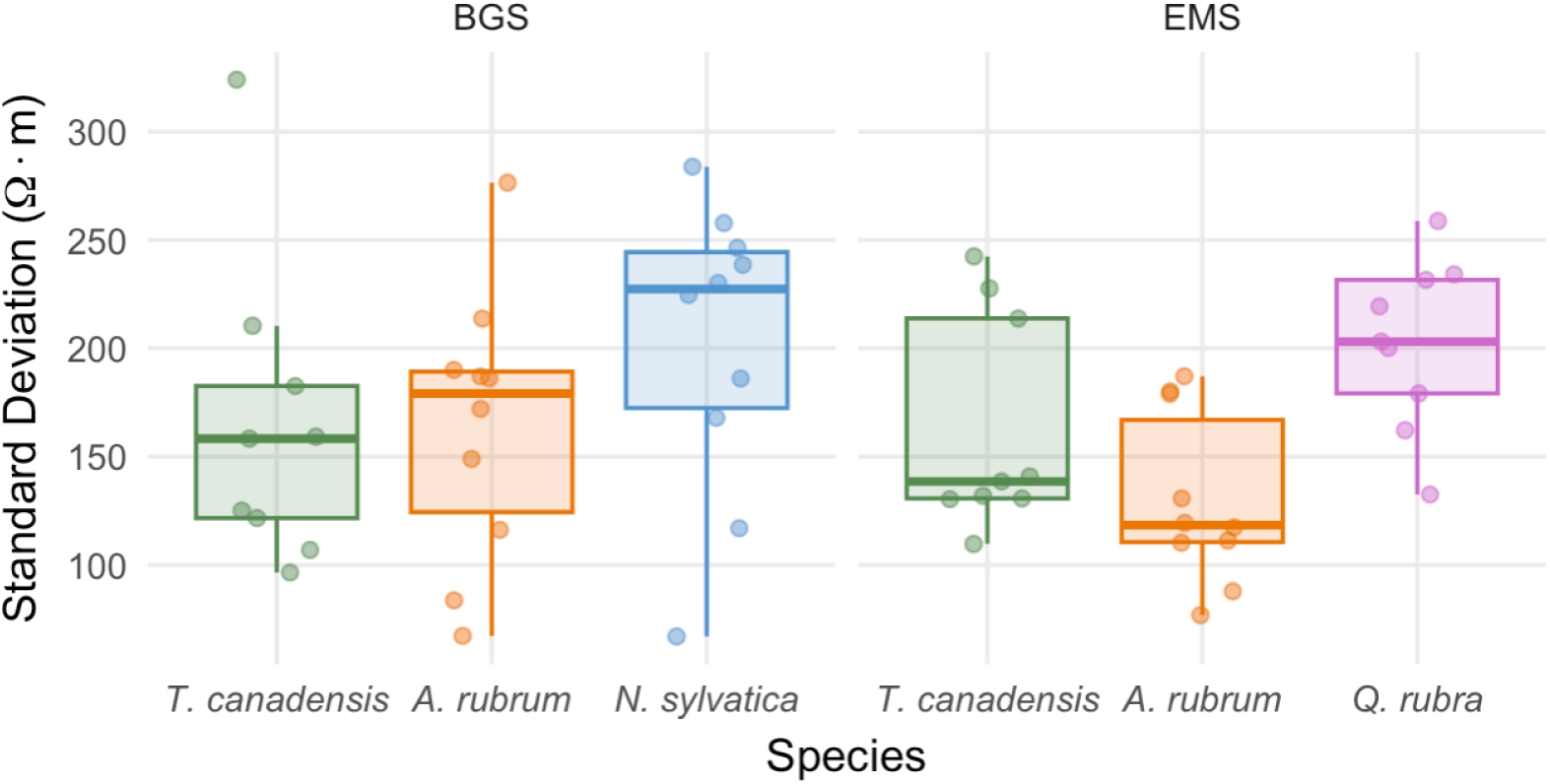
**Standard deviation (SD; Ω·m) of resistivity boxplots for each species within the BGS and EMS sites**. Jittered points represent raw data values computed by the open-source image analysis application. Higher SD values indicate more spatially heterogeneous resistivity distributions. There was no significant site effect (F(1, 51) = 1.08, p = 0.305) or species-by-site interaction (F(1, 51) = 0.77, p = 0.385), but there was a significant species effect (F(3, 51) = 3.23, p = 0.030). Averaged across sites, SD was marginally lower in A. rubrum (mean ± SE = 147 ± 12.0 Ω·m) than Q. rubra (202 ± 13.1 Ω·m, p = 0.082) and N. sylvatica (202 ± 21.4 Ω·m, p = 0.070). All other pairwise comparisons were not significant (p > 0.32).

**Figure S6.**
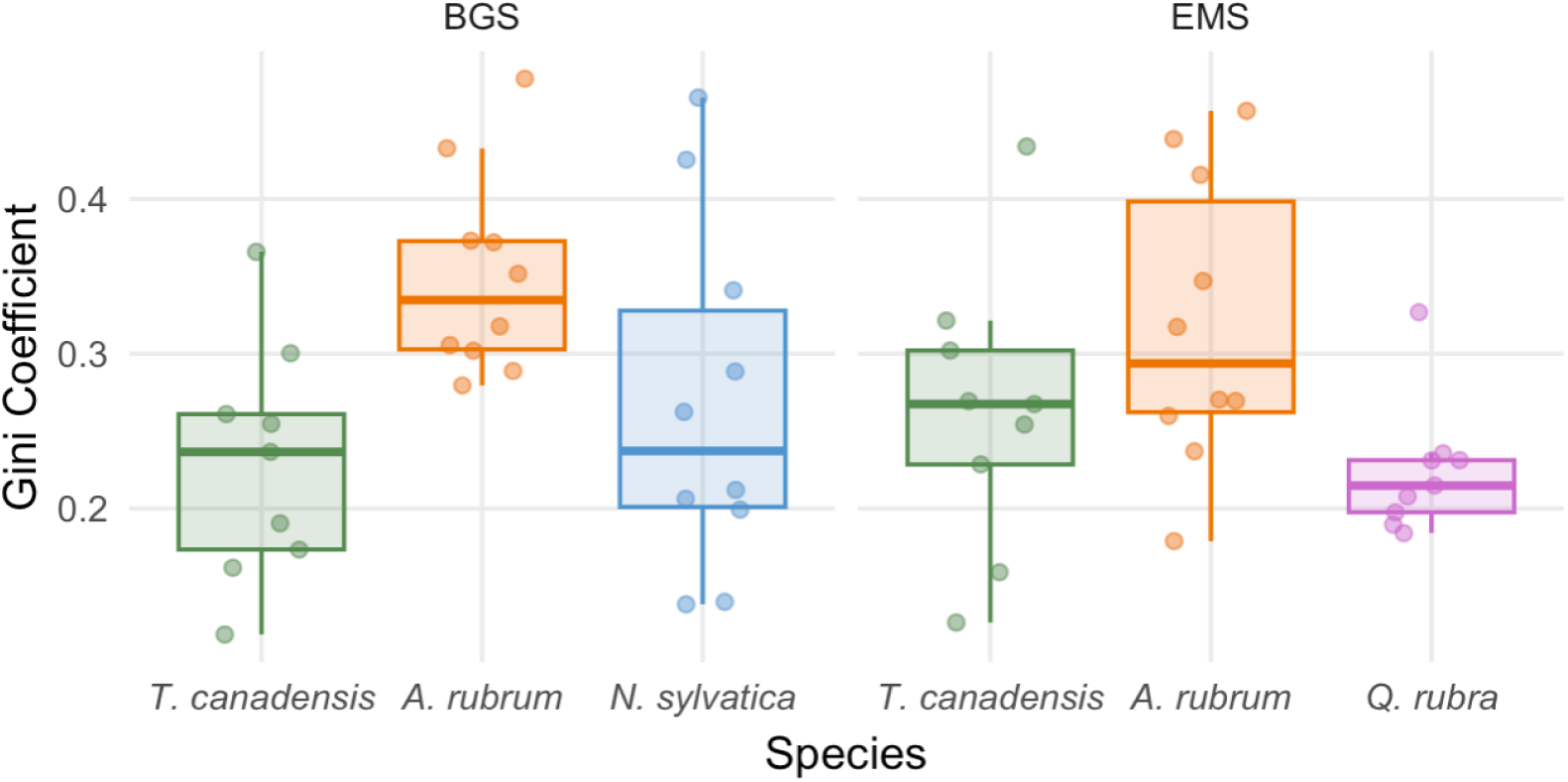
**Gini coefficient boxplots for each species within the BGS and EMS sites**. Jittered points represent raw data values computed by the open-source image analysis application. The Gini coefficient measures inequality in the resistivity distribution, ranging from 0 (uniform resistivity) to 1 (maximum inequality); higher values indicate greater concentration of extreme resistivity values. There was no significant site effect (F(1, 51) = 0.00, p = 0.98) or species-by-site interaction (F(1, 51) = 1.39, p = 0.24), but there was a significant species effect (F(3, 51) = 5.22, p = 0.003). Averaged across sites, the Gini coefficient was significantly higher in A. rubrum (mean ± SE = 0.34 ± 0.02) than T. canadensis (0.25 ± 0.02, p = 0.010) and Q. rubra (0.22 ± 0.01, p = 0.010). All other pairwise comparisons were not significant (p > 0.18).

**Figure S7.**
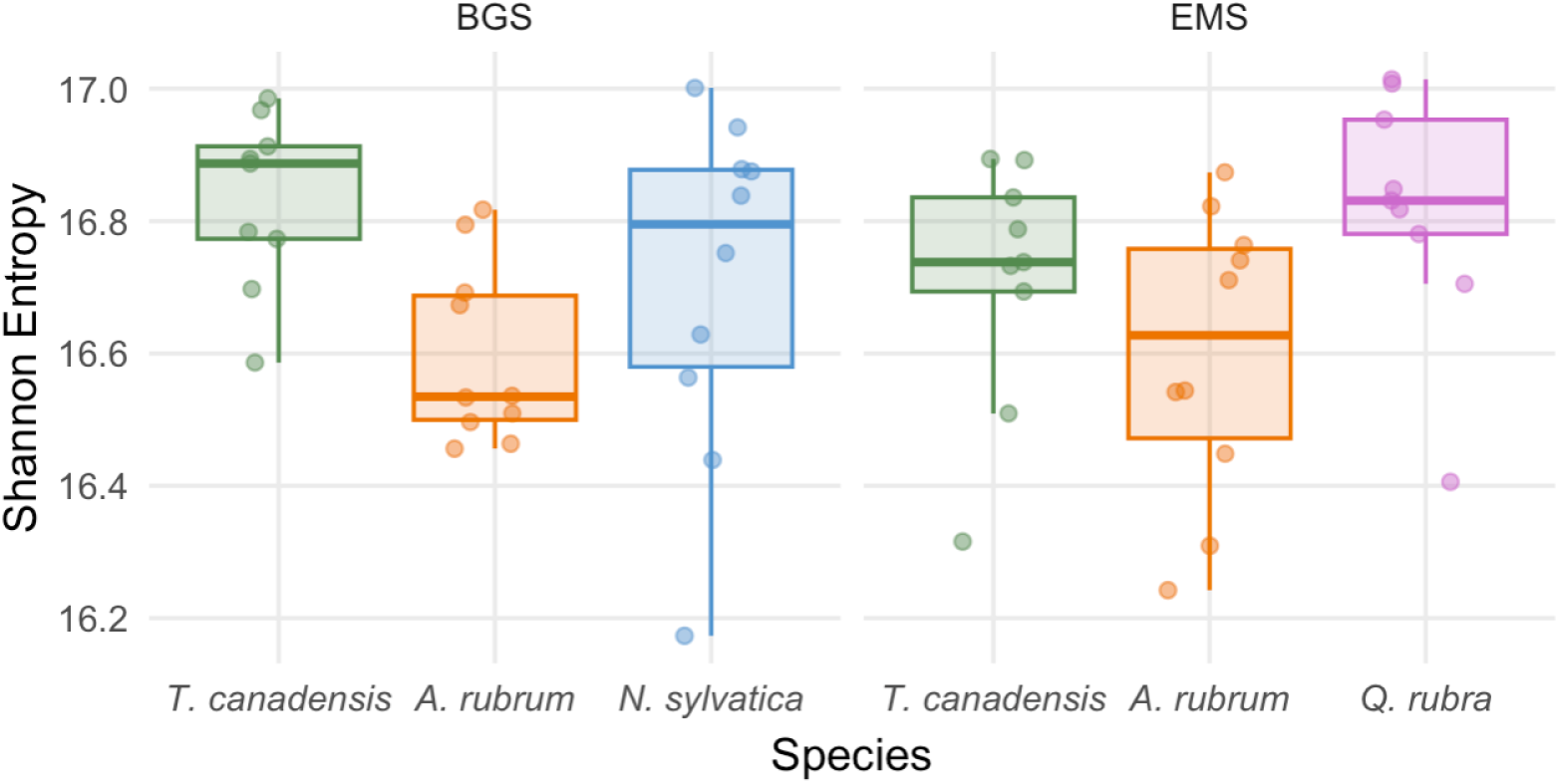
Shannon entropy boxplots for each species within the BGS and EMS sites. Jittered points represent raw data values computed by the open-source image analysis application. Shannon entropy measures heterogeneity in the resistivity distribution; higher values indicate more uniformly dispersed moisture patterns, while lower values indicate more concentrated or bimodal distributions. There was no significant site effect (F(1, 51) = 0.80, p = 0.375) or species-by-site interaction (F(1, 51) = 0.97, p = 0.329), but there was a significant species effect (F(3, 51) = 3.78, p = 0.016). Averaged across sites, Shannon entropy was significantly lower in A. rubrum (mean ± SE = 16.6 ± 0.04) than T. canadensis (16.8 ± 0.04, p = 0.039) and Q. rubra (16.8 ± 0.06, p = 0.032). All other pairwise comparisons were not significant (p > 0.45).

**Figure S8.**
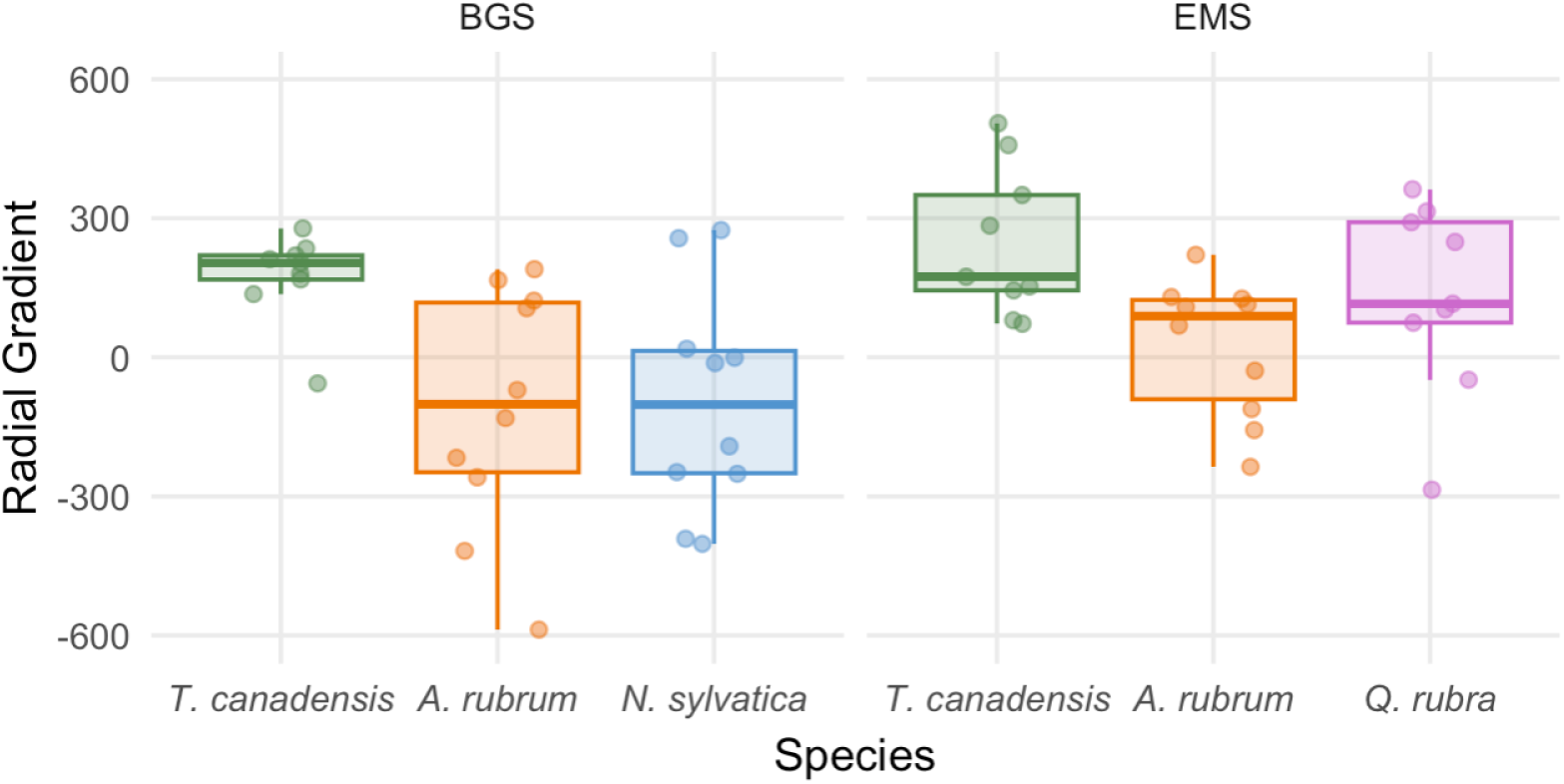
Radial gradient (Ω·m) boxplots for each species within the BGS and EMS sites. Jittered points represent raw data values computed by the open-source image analysis application. The radial gradient is the difference in mean resistivity between the outer and inner thirds of the cross-section. Positive values indicate drier edges relative to the center, typical of healthy sapwood; negative values indicate wetter centers relative to the edges, consistent with heartwood moisture accumulation or decay. There was no significant site effect (F(1, 51) = 2.67, p = 0.108) or species-by-site interaction (F(1, 51) = 0.24, p = 0.629), but there was a significant species effect (F(3, 51) = 7.83, p < 0.001). Averaged across sites, radial gradient was significantly greater in T. canadensis (mean ± SE = 211 ± 31.4 Ω·m) than A. rubrum (−43.1 ± 49.0 Ω·m, p = 0.001) and N. sylvatica (−94.7 ± 76.6 Ω·m, p = 0.001). The radial gradient of Q. rubra (131 ± 68.4 Ω·m) was marginally greater than N. sylvatica (p = 0.072). All other pairwise comparisons were not significant (p > 0.13).

**Figure S9.**
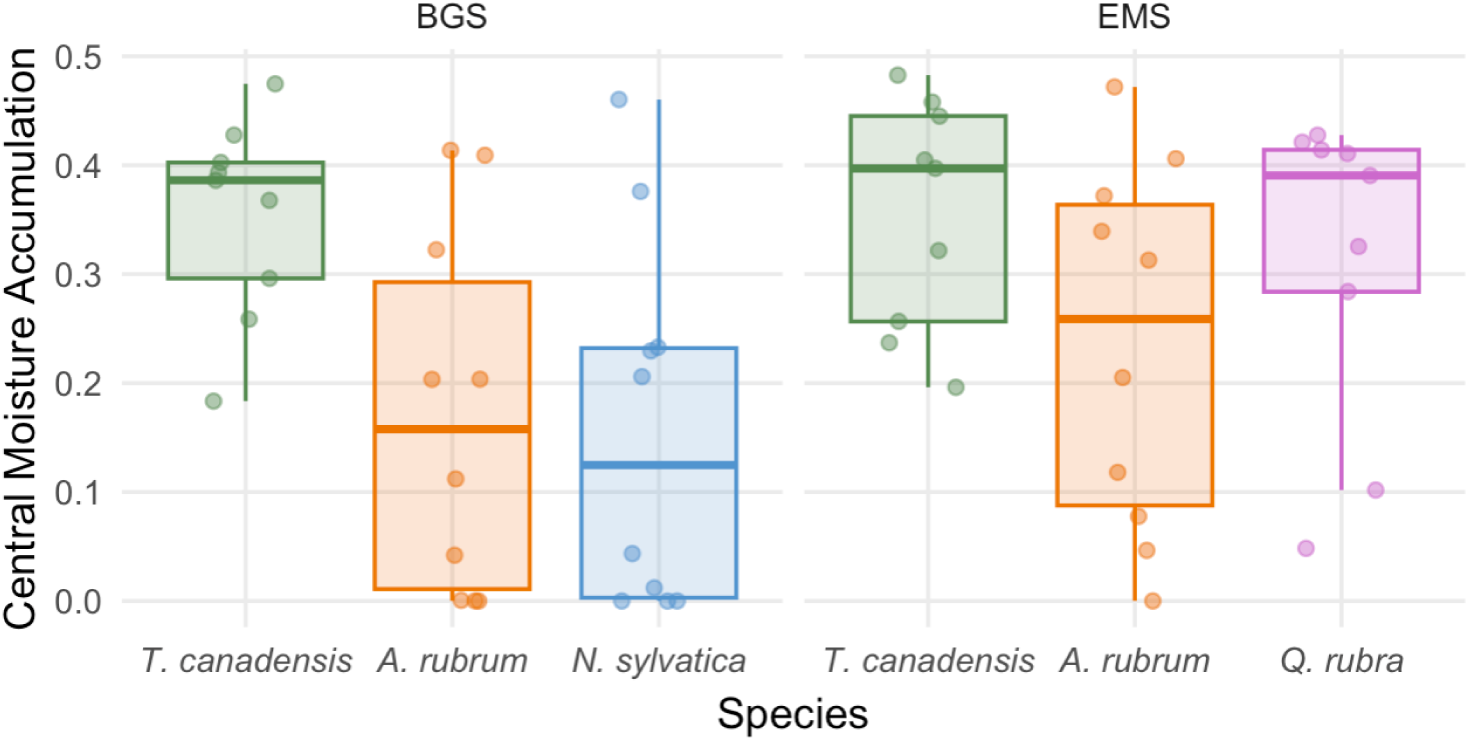
Central moisture accumulation (CMA) boxplots for each species within the BGS and EMS sites. Jittered points represent raw data values computed by the open-source image analysis application. CMA is the proportion of low-resistivity pixels (≤30th percentile of the cross-section) located in the inner third of the cross-section; values near 0.33 indicate proportional distribution throughout the cross-section, while values substantially above 0.33 indicate anomalous moisture concentration in the heartwood. There was no significant site effect (F(1, 51) = 0.525, p = 0.472) or species-by-site interaction (F(1, 51) = 0.45, p = 0.507), but there was a significant species effect (F(3, 51) = 5.68, p = 0.002). Averaged across sites, CMA was significantly higher in T. canadensis (mean ± SE = 0.36 ± 0.02) than A. rubrum (0.20 ± 0.04, p = 0.012) and N. sylvatica (0.16 ± 0.05, p = 0.006). All other pairwise comparisons were not significant (p > 0.09).

**Figure S10.**
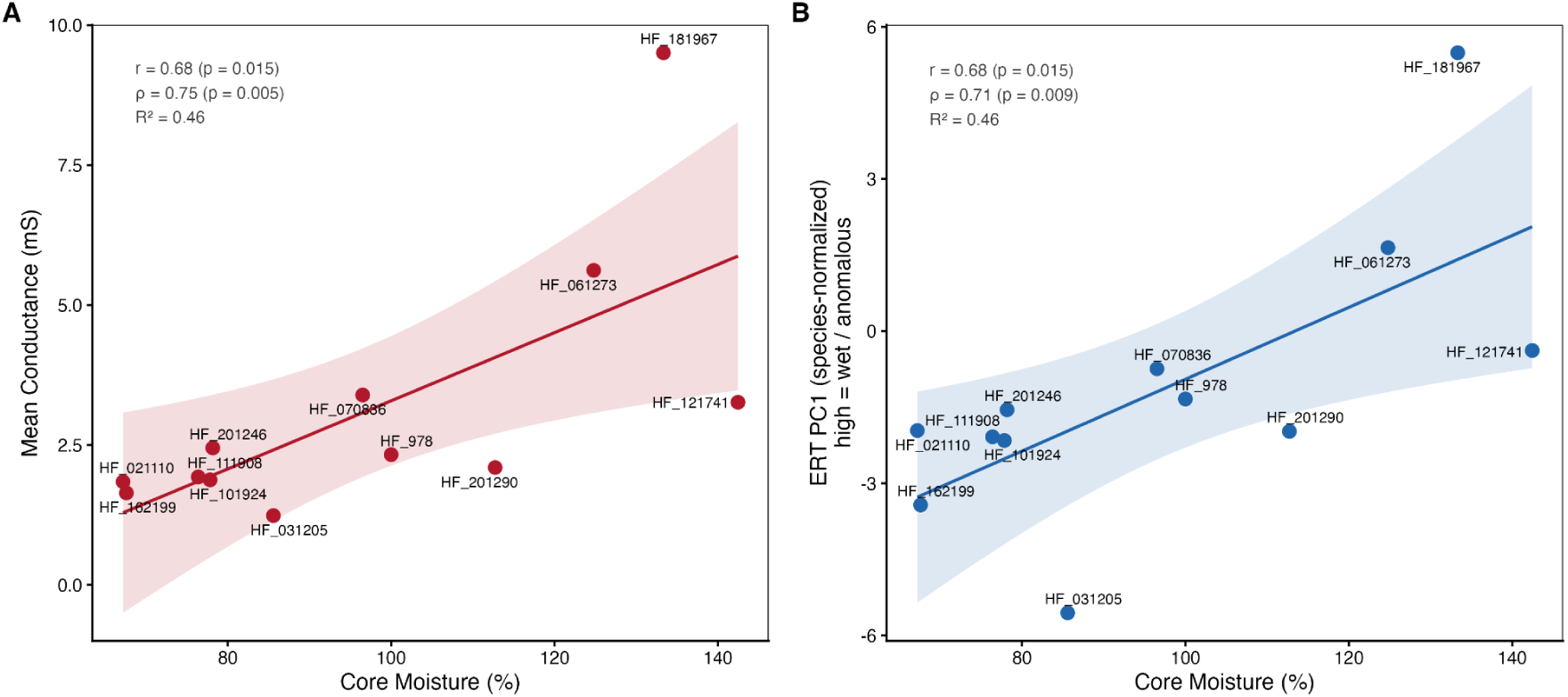
Relationship between gravimetric core moisture content and ERT-derived metrics for 12 *Tsuga canadensis* individuals. (A) Mean electrical conductance (mS; inverse of resistivity), a direct measure of wood moisture content. (B) ERT PC1 (species-normalized composite moisture anomaly index), capturing both moisture magnitude and spatial heterogeneity. Lines show ordinary least squares fits with 95% confidence bands. Pearson and Spearman correlation coefficients and R² are annotated in each panel.

## Notes

### Competing Interest Statement

The authors have declared no competing interest.

